# A universal platform for simultaneous TCRα/β removal enables safer and more potent TCR therapies and autoimmune modeling

**DOI:** 10.64898/2026.02.19.706929

**Authors:** Giorgia Zanetti, Mateusz Legut, Austin Chen, Farshid Fathi, Nathan Suek, Nato Teteloshvili, Hao Wei Li, Xiaolan Ding, Daniel Traum, Klaus H. Kaestner, Ryan E. Hoang, Edwin Bremer, Andrew K. Sewell, Audrey V. Parent, Remi J. Creusot, Megan Sykes, Mohsen Khosravi-Maharlooei

## Abstract

Adoptive T-cell therapies using tumour-specific T-cell receptors (TCRs) are limited by competition with endogenous receptors, which impairs efficacy and poses risks of off-target autoreactivity. Here we present a CRISPR-based platform that completely and selectively eliminates both endogenous TCR-α and -β chains without affecting introduced transgenic TCRs, irrespective of codon optimization. This approach achieves >90% deletion efficiency in Jurkat and primary human T cells, markedly enhancing the expression, pairing fidelity, and functional potency of transgenic receptors. Using a clinically relevant HLA-A*02:01-restricted DMF5 TCR, we show that dual TCR ablation boosts antigen-specific activation and cytotoxicity *in vitro* and significantly enhances tumor clearance *in vivo* in human immune system (HIS) mice, while preventing graft-versus-host disease (GVHD). Targeted locus amplification revealed that CRISPR-induced double-strand breaks did not alter lentiviral integration profiles, confirming genomic safety. Extending this approach to four insulin-reactive TCRs demonstrated that removal of endogenous receptors increased transduction efficiency and functional activity, with one (1E6) showing selective activation and infiltration of stem cell–derived islet grafts (SC-islets) *in vivo*. This study establishes a universal, safe, and scalable genome-editing platform for generating functionally precise human T cells. By integrating cancer immunotherapy and autoimmune disease modelling within a single framework, it provides a strong preclinical rationale for dual endogenous TCR removal as a route to improved specificity, safety, and therapeutic efficacy in TCR-based cell therapies.

## Introduction

TCR-based T cell therapy provides an alternative to chimeric antigen receptor (CAR) T cell therapy, by targeting intracellular antigens through T cell receptors, allowing for a broader range of cancer targets^1-5^. TCR-based adoptive T cell therapy emerged as one of the most promising approaches in cancer immunotherapy. However, the widespread clinical adoption of T cell therapies remains constrained by their reliance on patient-derived T cells and significant interpatient variability^6, 7^. In particular, individuals who have undergone chemotherapy and/or stem cell transplantation often experience, lymphopenia resulting in insufficient T cell numbers and impaired T cell function^8-11^. While allogeneic T cells address some limitations of autologous patient-derived T cells, they face poor persistence in allogenic recipients and can induce GvHD due to the lack of tolerance to recipient peptide-MHC complexes. The latter challenge can be addressed by genetically retargeting T cells through introduction of transgenic TCRs and simultaneously removing the endogenous TCRs to mitigate the risk of GvHD^12, 13^.

In contrast to CARs, TCRs recognize antigens derived from both extracellular and intracellular proteins presented in the context of the human leukocyte antigens (HLA)^14^. This broadens the spectrum of available antigens and enhances tumor specificity by targeting neo-antigens and other tumor-associated antigens^15^. Yet, the presence of endogenous TCR-α and -β chains in overall T cell populations impedes the correct expression of transgenic TCRs, due to mixed TCR dimer formation and competition for binding to the CD3 signaling complex^16, 17^. These mispaired TCRs not only reduce surface expression of the intended transgenic TCR but may also pose safety risks by generating unintended, and unknown, specificities that could lead to autoimmunity^18, 19^. The TCR-CD3 complex assembles in the endoplasmic reticulum before surface expression^20^, creating competition between endogenous and transgenic TCRs for CD3 association^21, 22^.

Several strategies have been explored to overcome TCR competition and mispairing, including of affinity-enhanced TCRs^23^, engineering mutations to improve pairing of transgenic TCRs^24^, and overexpression of CD3 components^21^. However, complete elimination of mispairing and restoration of optimal transgenic TCR expression has only been achieved via the full removal of endogenous TCRs^25-27^. Legut *et al*. reduced endogenous TCR expression by targeting the β-chain constant region with CRISPR while sparing codon-optimized transgenic TCRs^25^. However, this approach did not target TCR-α chain, and is not applicable to non-codon-optimized transgenic TCR constructs. Roth and colleagues instead inserted a new TCR construct within the constant region of TCR-α using homologous recombination, leaving the β-chain untouched^28^. Although adeno-associated virus (AAV), a commonly used gene-delivery vector, can deliver DNA repair templates for homology-directed repair (HDR) with high targeting efficiency^29^, non-viral single- or double-stranded DNA templates have not yet outperformed lentiviral transduction^28, 30^. Moreover, these AAV or non-viral systems necessitate complex sequence alignments with the endogenous TCR elements, making adaptation of existing viral vectors more challenging. More recently, Peters *et al*. employed CRISPR-Cas9 to delete the endogenous TCR-α chain to improve pairing of an exogenously introduced Type 1 diabetes (T1D)-associated TCR^31^. Importantly, their transgenic TCR construct included specific mutations to protect the introduced TCR from Cas9 cleavage improving on-target efficacy while reducing mispairing. However, the endogenous TCR-β remained intact, leaving residual potential for mispairing endogenous β- and introduced α-chains.

Here, we developed a novel CRISPR-based strategy that efficiently removes both endogenous TCR-α and -β chains without affecting the expression of transgenic TCRs, irrespective of codon-optimization. Using the clinically relevant DMF5 TCR targeting the melanoma antigen recognized by T cells-1 (MART-1) ^32^, we show that the simultaneous removal of both endogenous TCR-α and -β chains enhances the performance of transgenic T cells in a preclinical model of human metastatic melanoma. Importantly, extending this approach to autoimmune TCRs revealed that endogenous TCR removal also enhances transduction efficiency and functional activation, despite the generally low affinity of autoimmune TCRs^33^. Collectively, these findings establish a broadly applicable platform for both cancer and autoimmune TCR-based therapies.

## Results

### Design and validation of CRISPR strategy for exclusive removal of endogenous TCRs

To selectively remove endogenous TCRs without affecting our transgenic TCRs, whether they are codon-optimized or not, we designed guide RNAs (gRNAs) to target the intron-exon boundaries in the constant regions of both TCR-α and -β chains (Table S1). Since the transgenic TCR constructs lack introns, this design ensured that only endogenous TCR genes would be targeted. Several gRNAs directed against TCR-α constant region (*TRAC*), and the two TCR-β constant regions (*TRBC1* and *TRBC2*) were screened (Figure 1A). The *TRBC*-targeting gRNAs were designed against sequences conserved between *TRBC1* and *TRBC2*, enabling deletion of receptors containing either constant region. The Cas9 ribonucleoprotein (RNP) complexes containing different gRNA combinations were delivered into the Jurkat cells via electroporation. Four gRNAs, two for *TRAC* and two for *TRBC* were selected based on disruption efficiency. Dual targeting of each chain produced greater removal than individual gRNAs, and the combined use of all four guides achieved the highest level of endogenous TCR removal (more than 95%) (Figure 1B). The deletion results for non-efficient guides are shown in Figure S1.

**Figure 1.**
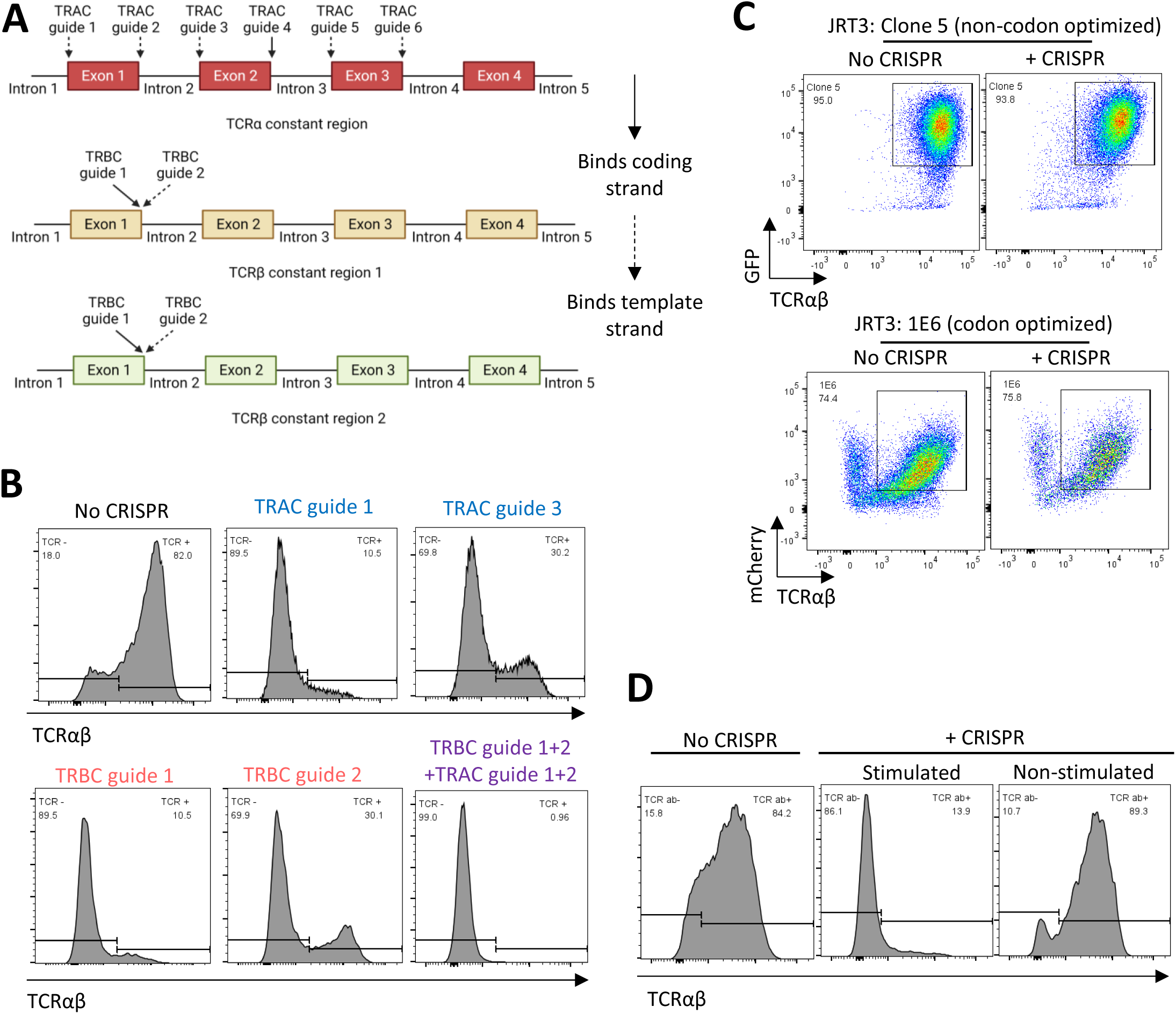
CRISPR strategy for selective removal of endogenous TCRs while preserving transgenic TCR expression. (A) Schematic representation of CRISPR strategy for removal of TCR-α and -β chains. Guide RNAs (gRNAs) were designed to target the TCR intron-exon junctions within the constant regions of *TRAC* and *TRBC* enabling efficient disruption of endogenous TCR expression while sparing transgenic receptors. (B) Surface expression of TCR in Jurkat cells electroporated with CRISPR-Cas9 with different gRNAs. (C) Surface TCR expression in JRT3-T3.5 (TCRβ^−/−^) cells electroporated with CRISPR-Cas9 using *TRAC* guide 1 and 3, and *TRBC* guide 1 and 2. Cells were transduced with either a non-codon optimized TCR (Clone 5) or codon-optimized TCR (1E6) (D) Surface expression of TCR in proliferating and resting PBMCs ± CRISPR.

To confirm that the gRNAs do not target transgenic TCRs, we tested our strategy on Jurkat, JRT3-T3.5 (TCRβ^−/−^) cells that were previously transduced with two different exogenous TCRs, one codon-optimized (1E6) and the other non-codon-optimized (Clone 5). Neither construct was affected by CRISPR editing, demonstrating that our gRNAs exclusively target endogenous sequences (Figure 1C). These results demonstrate that our CRISPR approach is compatible with existing TCR delivery platforms.

Because transcriptional activity and chromatin accessibility influence Cas9 efficiency ^34^, we next evaluated editing in primary human T cells. Peripheral blood mononuclear cells (PBMCs) were activated for 48 hours with anti-CD3/CD28/CD2–coated beads prior to electroporation with Cas9 RNPs containing the four gRNAs targeting *TRAC* and *TRBC*. Dual targeting of both the α and β chains achieved robust ablation in activated T cells, as assessed by flow cytometry analysis (Figure 1D). In contrast, minimal editing was observed in resting PBMCs, confirming that pre-activation is required for efficient CRISPR/Cas9-mediated disruption of TCR genes (Figure 1D).

### CRISPR removal of endogenous TCRs in primary human T cells enhances exogenous TCR expression

To optimize the timing and sequence of CRISPR-mediated removal of endogenous TCRs and lentiviral delivery of a transgenic TCR, we selected a clinically relevant TCR (DMF5) recognizing the melanoma antigen MART-1, which is widely overexpressed in melanomas^35^. A codon-optimized MART-1-specific DMF5 TCR restricted by HLA-A*02:01 was used. We compared three strategies for combining CRISPR editing and lentiviral transduction: (i) lentiviral transduction followed by CRISPR (V then C), (ii) simultaneous CRISPR and lentiviral delivery C+V), and (iii) CRISPR editing followed by lentiviral transduction (C then V) (Figure 2A). Lentiviral transduction followed by CRISPR did not enhance transduction efficiency relative to lentivirus-only controls (V) and resulted in marked cell death and delayed expansion (Figure S2A). These cells in the V then C group exhibited significant cell death and delayed expansion (Figure S2B) (V stands for virus and C stands for CRISPR). In contrast, both simultaneous (C+V) and sequential (C then V) editing-transduction approaches significantly increased transduction efficiency compared to the V group (Figure 2B and S2A).

**Figure 2.**
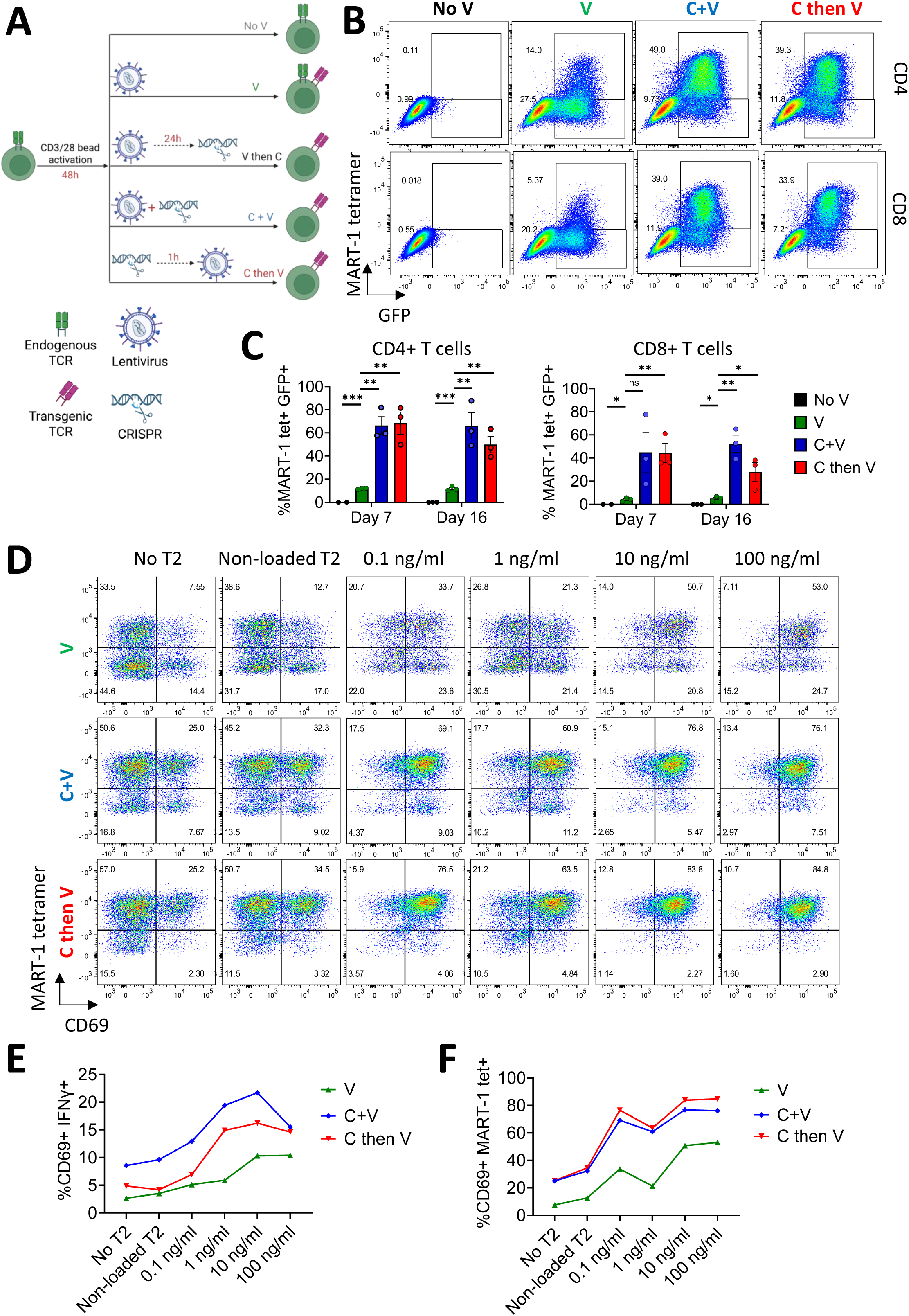
CRISPR-mediate removal of endogenous TCRs enhances transgenic DMF5 TCR expression and function in primary human T cells. (A) Schematic presentation of optimization of timing and order of DMF5 TCR transduction and CRISPR removal of endogenous TCR-α and TCR-β chains. (B) Expression of DMF5 TCR (tetramer staining) and reporter gene (GFP) in primary T-cells following different conditions of lentiviral transduction of DMF5 TCR with and without CRISPR-removal of endogenous TCRs: no lentiviral transduction (No V), lentiviral transduction only (V), simultaneous CRISPR-removal of endogenous TCRs and lentiviral transduction (C+V), and CRISPR followed by lentiviral transduction (C then V). (C) Expression of GFP and DMF5 TCR in primary T cells at day 7 and day 16 post-transduction. Data represents the mean±SEM from three independent experiments. Statistical significance was determined by one-way ANOVA (* 0.01<p-value<0.05, **0.001<p-value<0.01, ***p-value<0.001). (D) Representative flow plots showing antigen-specific activation of DMF5 TCR-transduced T cells, in V, C+V and C then V groups, upon recognition of MART-1 peptide presented on HLA-A2^+^ T2 cells as indicated by CD69 surface upregulation. T2 target cells were pulsed with graded concentrations of MART-1 peptide and co-cultured with transduced T cells for 24 h. (E) Percentage of CD69^+^IFNɣ^+^ T cells following stimulation with T2 cells loaded with different concentrations of MART-1 peptide. (F) Percentage of CD69 and MART-1 tetramer double positive T cells in response to T2 cells loaded with increasing concentrations of MART-1 peptide.

In both C+V and C then V conditions, the transgenic DMF5 TCR was stably expressed through *ex vivo* expansion (days 7-16; Figure 2C). The comparable proliferation of transduced and non-transduced T cells lacking endogenous TCRs indicates that CRISPR-mediated removal of the native receptor does not impair T-cell survival or homeostatic expansion in the presence of IL-2, IL-7, and IL-15, even in the absence of sustained TCR signaling (Figure 2C).

Across independent experiments, the ratio of MART-1 tetramer^+^ to tetramer^neg^ cells within GFP^+^ (transduced) T cells ranged from 0.5 to 2 in the V group (Figure 2B and S2C), suggesting that a substantial proportion of transduced T cells expressed mispaired or incomplete TCRs. In contrast, this ratio increased to 4-10 in the C+V and C then V groups, demonstrating that elimination of endogenous TCRs markedly enhances correct pairing and surface expression of the intended transgenic TCR while reducing the risk of generating hybrid receptors with novel, potentially autoreactive specificities (Figure 2B and S2C).

### TCR-transduced T cells lacking endogenous TCRs show enhanced activation upon antigen recognition

To evaluate antigen-specific activation, DMF5 TCR-transduced T cells from the V, C+V, and C then V groups were co-cultured with HLA-A2^+^ T2 cells pulsed with graded concentrations of MART-1 mimotope peptide (26-35, ELAGIGILTV). The human-derived lymphoid cell line T2 lacks TAP gene expression and is impaired in presenting endogenous peptides^36^. After 24 hours of co-culture, the expression of activation markers CD69 and IFN-ɣ increased significantly in a peptide dose-dependent manner across all three conditions. Notably, this activation was substantially greater in the C+V and C then V groups, in which endogenous TCRs were removed, compared to the V group (Figure 2D, E and F). These findings are consistent with the higher frequency of MART-1 tetramer^+^ cells in the C+V and C then V groups (Figure 2B), demonstrating that elimination of endogenous TCRs enhances both the surface expression and antigen-dependent activation of transgenic TCRs.

### Eliminating endogenous TCRs enhances the ability of TCR-transduced T cells to kill autologous tumor cells expressing cognate antigens

To establish an autologous model for testing antigen-specific cytotoxicity, PBMCs were obtained from a healthy HLA-A02:01^+^ donor. CD19^+^ B cells were magnetically sorted and transformed with Epstein-Barr virus (EBV) to generate a B-lymphoblastoid cell line (LCL), while the remaining PBMCs were cryopreserved subsequent assays. A subset of LCLs were transduced with a lentiviral vector encoding the MART-1 peptide (H3 vector). Because lentiviral transduction efficiency varied between 3-18% under different conditions, H3-transduced (mCherry^+^) LCL cells were enriched by fluorescence-activated cell sorting (FACS) and expanded for downstream assays (Figure 3A). Despite sorting, the purity of mCherry^+^ cells remained approximately 50-60% and gradually decreased over time.

**Figure 3.**
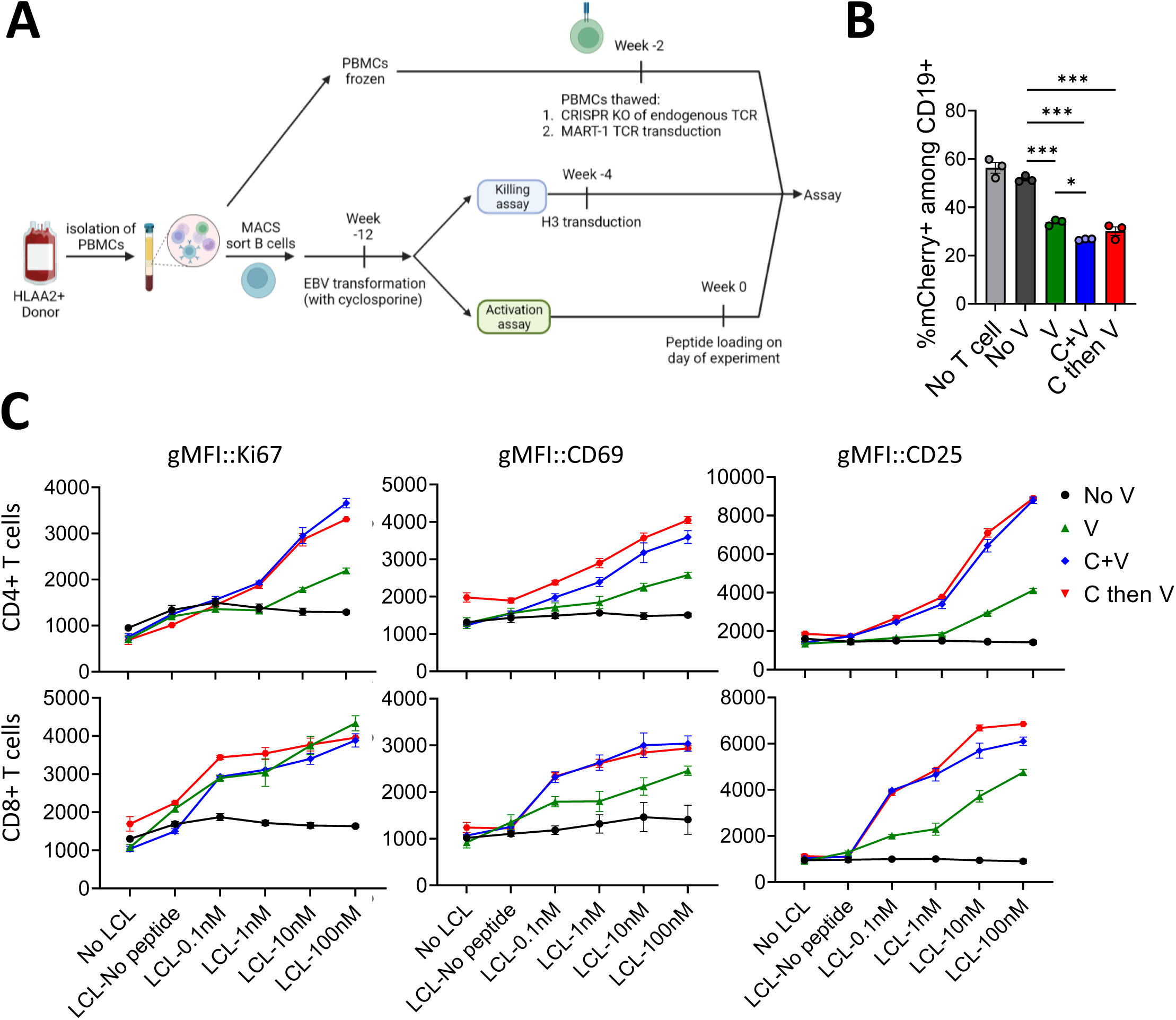
Endogenous TCR removal enhances activation and cytotoxicity of DMF5 TCR-transduced T cells against autologous target cells. (A) Schematic overview of the generation of autologous LCL cells engineered to express MART-1 peptide via the H3 construct, used as autologous antigen-presenting stimulators for DMF5 TCR-transduced T cells. (B) *In vitro* cytotoxicity assay showing the percentage of remaining mCherry^+^ autologous LCL cells expressing MART-1 peptide after 48 h co-culture with DMF5 TCR-transduced T cells (n=3 replicates per group). Data are presented as mean±SEM. Statistical significance was determined by one-way ANOVA (* 0.01<p-value<0.05, **0.001<p-value<0.01, ***p-value<0.001). (C) *In vitro* activation of DMF5 TCR-transduced T cells in response to autologous LCL cells loaded with different concentrations of MART-1 peptide. (V: virus, C: CRISPR)

Cryopreserved PBMCs from the same donor were thawed, activated with anti-CD3/CD28/CD2 beads for 48 hours, and transduced with a lentiviral vector carrying DMF5 TCR under different conditions with and without removal of endogenous TCRs. To assess antigen-specific cytotoxicity, DMF5 transduced T cells (V only, C+V and C then V), and untransduced controls (No V) were co-cultured for 48 hours with H3-transduced autologous LCLs. The frequency of mCherry^+^ target cells among total B cells was then quantified. T cells from both the C+V and C then V groups demonstrated the highest levels of antigen-specific cytotoxicity, with the C+V group showing the strongest cytotoxic activity relative to both the untransduced (No V) and lentivirus-only V group controls (Figure 3B).

Because the autologous LCL system was limited by variable transduction efficiency of LCLs and declining MART-1 peptide transgene expression, we next assessed T-cell function using peptide-loaded autologous LCLs. In line with our earlier observations, removal of endogenous TCRs (C+V and C then V groups) significantly enhanced T-cell proliferation (Ki-67) and activation (CD69 and CD25), particularly at higher peptide concentrations (Figure 3C). Together, these data demonstrate that CRISPR-mediated elimination of endogenous TCRs improves both the activation and antigen-specific cytotoxicity of TCR-engineered T cells in an autologous setting.

### TCR-transduced T cells lacking endogenous TCRs display enhanced *a*ctivation and specific cytotoxicity against HLA-A2^+^ MART-1-expressing K562 cells

Because of the limited transduction efficiency and unstable transgene expression in LCL cells, we explored an alternative tumor model that permits antigen-specific recognition while avoiding alloreactivity. The erythroleukemia cell line K562, which lacks HLA class I and II expression ^37^, was selected to minimize allogeneic responses from donor T cells. To confirm the ability of these cells to present MART-1 peptide to the HLA-A2-restricted DMF5 TCR, we utilized a K562 cell line that had been stably transduced with the HLA-A*0201 gene^38^ and further transduced it with a lentiviral construct encoding MART-1 peptide (H3 construct) (Figure S3A). As T cells from HLA-A2^+^ donors are tolerant to self HLA A2, this system provides a clean background for assessing MART-1–specific responses. To verify that MART-1–peptide– expressing K562 cells (K562-H3) can induce DMF5 TCR activation, we co-cultured them for 24 hours with TCR-KO Jurkat (JRT3-T3.5) cells transduced with the DMF5 TCR. DMF5 TCR⁺ JRT3-T3.5 cells upregulated CD69 upon co-culture with K562-H3, confirming TCR activation (Figure S3B).

Human T cells from an HLA-A2^+^ donor were transduced with the DMF5 TCR under different conditions (No V, V, C+V, and C then V) with or without removal of endogenous TCRs. After a 48 h of co-culture, with mCherry^+^ HLA-A2^+^ K562 cells, activation of DMF5 TCR–transduced T cells was assessed by expression of Ki-67, HLA-DR, and CD25. Only MART-1 peptide–expressing (mCherry⁺) HLA-A2⁺ K562 cells induced activation, confirming antigen dependence. While T cells in the V group showed moderate activation, the C+V and C then V groups displayed the highest expression of activation markers (Figure 4A). In contrast, untransduced T cells (No V) showed no response, confirming the absence of cross-reactivity through endogenous receptors. These results demonstrate that removal of endogenous TCRs enhances the antigen-specific activation of DMF5 TCR– transduced T cells.

**Figure 4.**
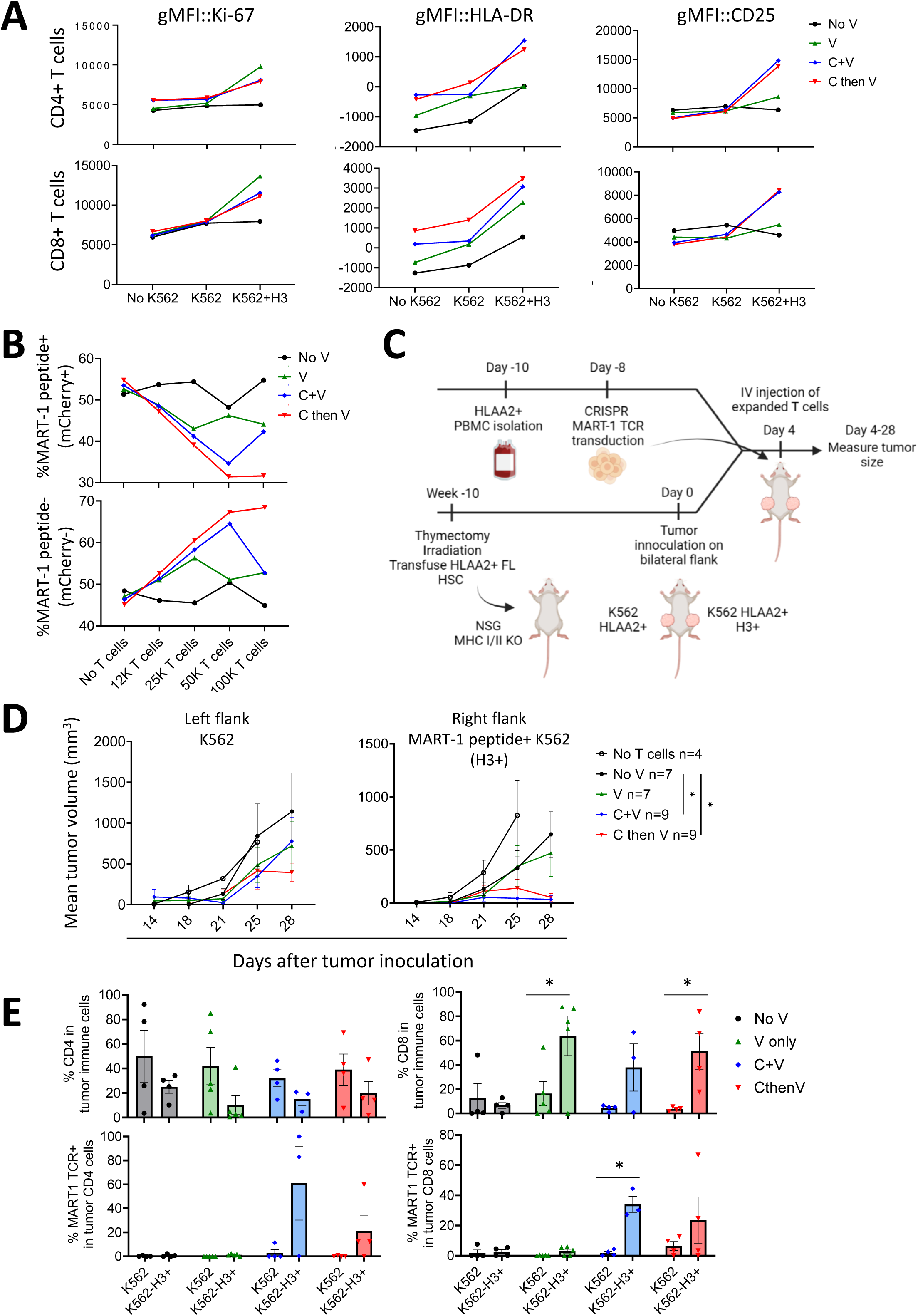
Endogenous TCR KO enhances the tumor-specific cytotoxicity of DMF5 TCR-transduced T cells *in vitro* and *in vivo*. (A) Proliferation (Ki-67) and activation (HLA-DR and CD25) of HLA_A*02:01^+^ DMF5 TCR-transduced T-cells after 48h of co-culture with mCherry^+^ HLA-A2^+^ K562 tumor cells, or unmodified HLA-A2^+^ K562 cells (negative control). (B) *In vitro* cytotoxicity of engineered T cells against a 1:1 mix of mCherry^+^ HLA-A2^+^ unmodified K562 and HLA-A2^+^ cells (100,000 total targets) after 48 hours. Cytotoxicity was measured across multiple effector to target ratios. (C) Schematic overview of the humanized xenograft tumor model. NSG MHC I/II double knockout reconstituted with a human immune system were intradermally implanted with mCherry^+^ HLA-A2^+^ K562 cells in their right flank and untransduced HLA-A2^+^ K562 cells in their left flank. T cells were adoptively transferred three days post-implantation, and tumor growth was monitored over several weeks. (D) Growth curves of mCherry^+^ HLA-A2^+^ K562 and HLA-A2^+^ K562 tumors over time (n=4-9 per group). Data represent mean ± SEM; two-way ANOVA was used for statistical analysis (* 0.01<p-value<0.05). (E) Frequency of infiltrating total and MART-1-specific T cells within mCherry^+^ HLA-A2^+^ K562 and untransduced HLA-A2^+^ K562 tumor grafts (n=4-per group). Data represent mean ± SEM, paired T test was used for statistical analysis (* 0.01<p-value<0.05). (V: virus, C: CRISPR).

To evaluate cytotoxic function, we co-cultured DMF5 TCR-transduced T cells with a 1:1 mixture of HLA-A2^+^ K562 cells expressing (mCherry^+^) or lacking (mCherry^-^) MART-1 peptide. After 48 hours, only T cells expressing DMF5 TCR selectively eliminated the mCherry^+^ target cells, sparing the mCherry^-^ controls (Figure 4B). The C+V and C then V groups exhibited greater cytotoxicity, which scaled with the effector-to-target ratio, confirming both the antigen specificity and functional superiority of TCR-transduced T cells lacking endogenous TCRs.

### Removal of endogenous TCRs enhances *in vivo* tumor cytotoxicity of TCR-transduced T cells

To evaluate the in vivo functionality of transgenic T cells, we employed a human immune system (HIS) xenograft tumor mouse model^39^, using NSG-MHC I/II double-knockout (DKO) mice. This strain combines the immunodeficient background of nonobese diabetic severe immunodeficient γ chain null (NSG) mice with MHC class I and II deficiency, which delays in the onset of GvHD following human PBMCs transfer^40^. Mice underwent thymectomy, received sublethal irradiation (1Gy), and were reconstituted with HLA-A02:01^+^ fetal liver-derived CD34^+^ hematopoietic stem cells (HSCs) to establish human antigen-presenting cells (APCs) in the absence of human T cells (Figure 4C and S5A). Ten weeks after reconstitution, mice were implanted subcutaneously mCherry^+^ HLA-A2^+^ K562 cells (expressing MART-1 peptide) in their right flank and untransduced (mCherry^-^) HLA-A2^+^ K562 cells in their left flank. Three days later, the mice received intravenous injections of 1x10^6^ DMF5 TCR-transduced T cells under different conditions (No V, V, C+V and C then V using an HLA-A2^+^ donor) or PBS as control (Figure 4C). Adoptive transfer of T cells from the C+V and C then V groups significantly suppressed the growth of mCherry^+^ HLA-A2^+^ K562 cells compared to T cells (No V) and V-only T-cell groups. Tumor suppression was antigen-specific, as no inhibition of untransduced HLA-A2^+^ K562 tumors was observed (Figure 4D).

Endpoint analyses revealed a preferential infiltration of CD8^+^ T cells into mCherry^+^ HLA-A2^+^ tumors in all groups receiving DMF5 TCR-transduced T cells (V, C+V, and C then V). However, robust infiltration of MART-1 tetramer^+^CD8^+^ T cells in mCherry^+^ tumors was observed only in the C+V and C then V groups, reaching statistical significant in the C+V group (Figure 4E). There was a significant inverse correlation between the number of HLA-A2^+^ mCherry^+^ K562 tumor cells and the number of MART-1 tetramer^+^ CD8^+^ infiltrating T cells present in tumors explanted from recipient HIS mice (Figure S4A).

Hematoxylin and eosin (H&E) staining of explanted tumors confirmed increased lymphocyte infiltration in mCherry^+^ HLA-A2^+^ K562 tumors from mice treated with of DMF5 TCR-transduced T cells, particularly in the C+V and C then V groups (Figure 5). Quantification of lymphocyte density within tumour sections further demonstrated that significant immune infiltration occurred only in mice receiving T cells lacking endogenous TCRs (Figure S4B). Together, these results show that removal of endogenous TCRs markedly enhances both the tumor infiltration and antigen-specific cytotoxicity of TCR-engineered T cells *in vivo*.

**Figure 5.**
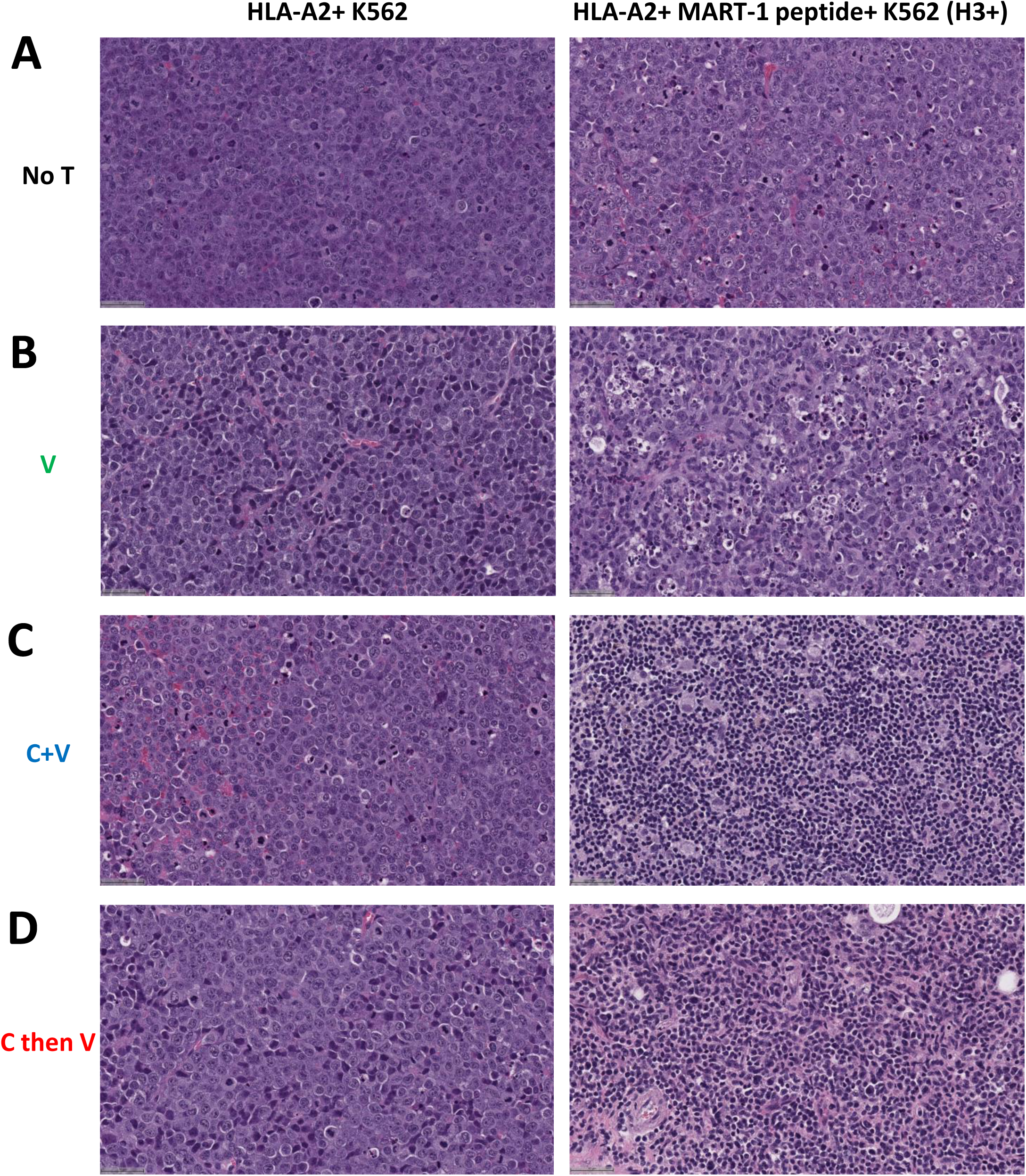
CRISPR-mediated removal of endogenous TCRs increases immune infiltration specifically in HLA-A2^+^ MART-1 peptide-expressing tumors. Representative histological images (H&E staining) of HLA-A2^+^ K562 and mCherry^+^ HLA-A2^+^ K562 tumors. (A) In mice that did not receive T cell transfer (No T), both tumor types showed minimal immune cell infiltration. (B) In mice receiving DMF5 TCR-transduced T cells without CRISPR editing (V), modest T cell infiltration was in mCherry^+^ HLA-A2^+^ K562 tumors compared to the control HLA-A2^+^ K562 tumors. (C) In mice receiving simultaneously CRISPR-edited and transduced T cells (C+V) and (D) sequentially edited and transduced T cells (C then V), mCherry^+^ HLA-A2^+^ K562 tumors exhibited markedly increased T-cell infiltration relative to control tumors (n=3 mice per group).

### CRISPR-mediated removal of endogenous TCRs prevents GvHD in HIS mice

To determine whether the elimination of endogenous TCRs mitigated off-target reactivity of TCR-transduced T cells, we assessed GvHD in HIS mice. NSG–MHC I/II DKO mice were thymectomized and reconstituted with human fetal liver-derived CD34^+^ HSCs, generating human APCs while excluding human T cells (Figure 6A and S5B). Human cell reconstitution in these mice are shown in Figure S5. Mice were infused with either untransduced (No V), lentivirus-only (V), or CRISPR-edited T cells (C+V or C then V). Among recipients of untransduced T cells, five out of six developed lethal GvHD within 46 days of transfer and all mice receiving V-group T cells (n=6) similarly succumbed to GvHD (Figure 6B-D). In contrast, none of the mice receiving C+V cells (n=5) developed disease, and only one recipient of the C then V T cells (n=6) died within the 46-day follow-up, likely unrelated to GvHD (Figure 6D).

**Figure 6.**
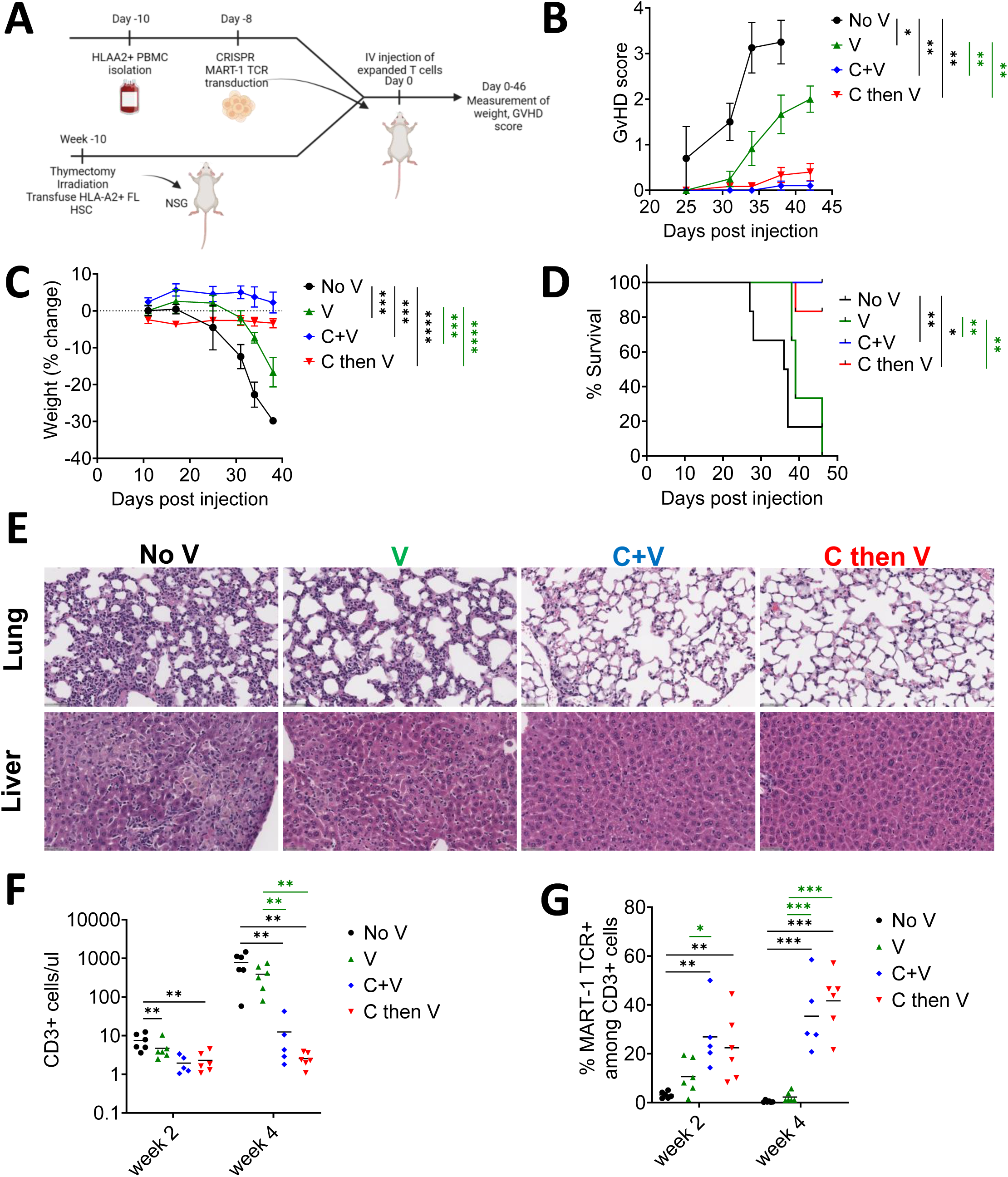
CRISPR removal of endogenous TCRs prevents GvHD following adoptive transfer of DMF5 TCR-transduced T cells. (A) Schematic overview of the experimental design used to assess GvHD in recipient HIS mice following adoptive transfer of DMF5 TCR-transduced T cells with and without CRISPR-mediated removal of endogenous TCRs. (B) GvHD scores and (C) percentage of weight loss in recipient mice after transfer of DMF5 TCR-transduced T cells prepared under different conditions (n=5-6 per group). (D) Survival curves of recipient mice receiving TCR-transduced T cells with or without endogenous TCR removal. (E) Representative H&E stained histological sections of lungs and the livers from recipient mice. Prominent lymphoid infiltrates were evident in mice that received unedited (no V) or lentivirally transduced (V) cells, but absent in mice given CRISPR-edited T-cells (C+V or C then V). (n=5-6 per group). (F) Absolute number of CD3^+^ T cells per µl of peripheral blood at 2 and 4 weeks after adoptive transfer. (G) Frequency of DMF5 TCR^+^ T cells in peripheral blood 2 and 4 weeks after adoptive transfer. Data in panels B and C were analyzed by two-way ANOVA and presented as mean ± SEM. Panels F and G were analyzed by one-way ANOVA. Survival data (D) were analyzed using the log-ranked (Mantel-Cox) test (* 0.01<p-value<0.05, **0.001<p-value<0.01, ***p-value<0.001, ****p-value<0.0001). Black asterisks indicate statistical differences between the “No V” group and other groups, while green asterisks indicate differences between the “V” group and other groups.

Clinical manifestations of GvHD including significant body weight loss and typical histopathological features (mononuclear infiltration in the liver and lung and bronchial narrowing) were observed exclusively in No V and V groups, but were absent in C+V and C then V mice (Figure 6B, C, and E). Peripheral analysis revealed marked expansion of human CD3^+^ T cells between weeks 2 to 4 post transfer only in the No V and V groups (Figure 6F). This expansion, driven by endogenous TCR recognition of mouse antigens, is characteristic of graft-versus-host (GvH)-reactive T cells. In contrast, T-cell numbers remained stable in C+V and C then V mice, consistent with the absence of GvHD. Moreover, while the proportion of MART1 tetramer^+^ T declined between weeks 2 to week 4 in the V group, this population remained stable in both CRISPR-edited groups (Figure 6G), confirming that removal of endogenous TCRs eliminates xenoreactive expansion while preserving the persistence of antigen-specific transgenic T cells.

### CRISPR-induced double-strand DNA breaks do not alter the integration profile of lentivirally-delivered transgenic TCRs

CRISPR-mediated editing generates double-strand DNA breaks (DSBs) at *TRAC* and *TRBC* loci raising the possibility that lentiviral transgenes could preferentially integrate into these regions as DSBs can “capture” exogenous DNA^41, 42^. We therefore hypothesized that the integration profile of the DMF5 TCR transgene might differ between the of C+V versus C then V groups. To test this, we performed Targeted Locus Amplification (TLA) sequencing to map precise transgene integration sites in the human genome, and their genomic context. Two primer sets targeting the RRE or GFP regions of the DMF5 TCR transgene were designed for the TLA analysis (Table S2). Genomic DNA from viable cryopreserved T cells of the C then V and C+V groups was processed using the Cergentis protocol^43^ and sequencing reads were aligned to the human reference genome (GRCh38/hg38). Sequence coverage plots showed no large coverage peaks in either samples, indicating that no genomic region was disproportionately targeted by lentiviral integration. Importantly, no integration breakpoints were detected within the *TRAC* (chr14:22,547,506-22,552,156) or *TRBC* (chr7:142,299,011-142,813,287) loci, excluding preferential integration at CRISPR-induced DSBs (Figure 7A and 7C).

**Figure 7.**
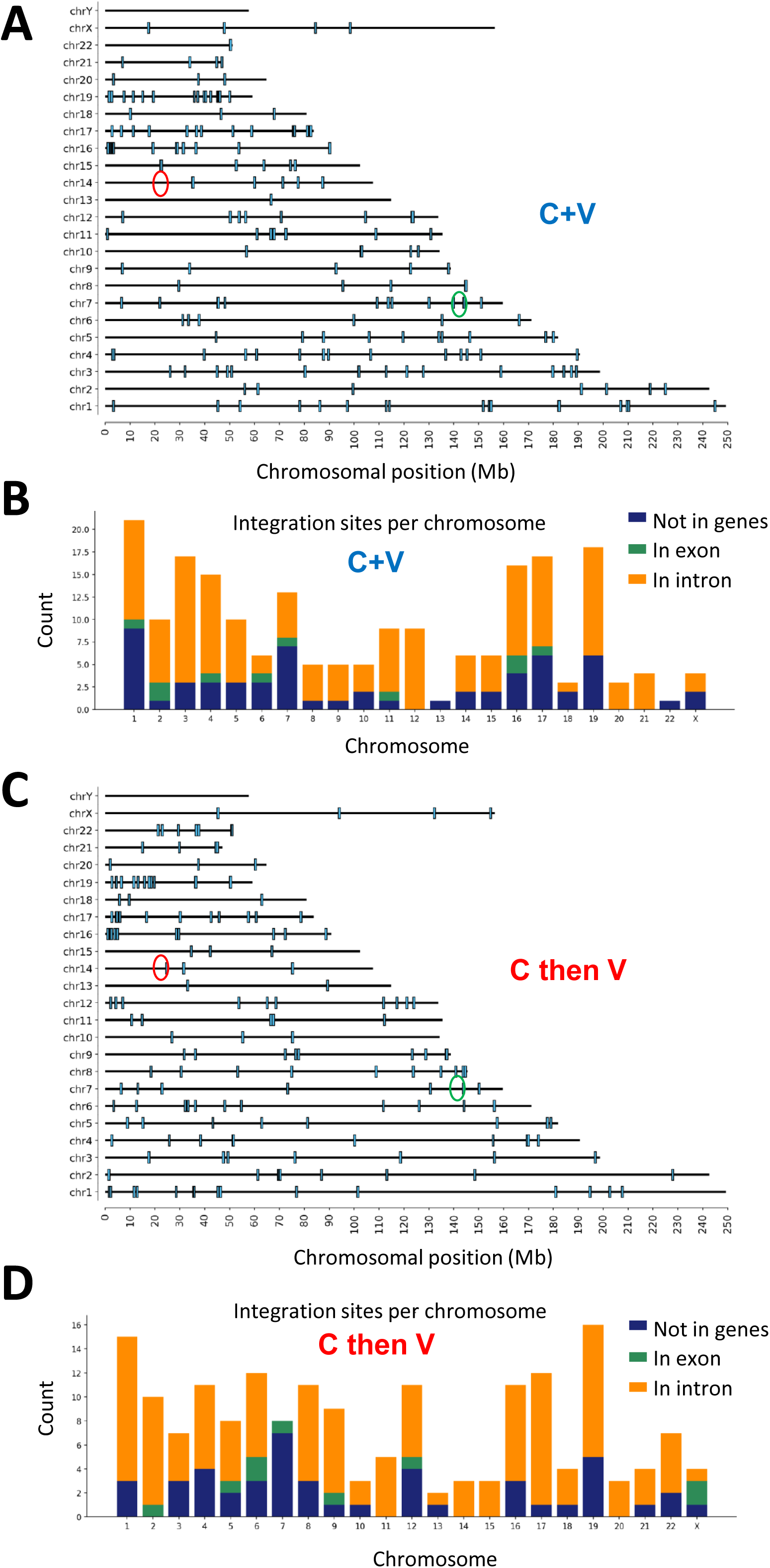
Targeted Locus Amplification (TLA) reveals widespread but safe integration of the DMF5 TCR transgene. (A) and (C) Genome-wide distribution and characterization of the integration sites of virally delivered DMF5 TCR transgene in human primary T cells edited under C+V and C then V conditions, respectively. Integration sites were equally distributed across all chromosomes, with no insertions detected in the *TRAC* (red circle) or *TRBC* (green circle) loci. (B) In DMF5 TCR-transduced T cells from C+V group, 179 total integration sites were identified, 9 (5%) within gene exons, 124 (69%) within gene introns and 2 (1%) within 1 kb upstream of annotated genes. (D) In DMF5 TCR-transduced T cells from C then V group, 204 integration sites were detected, 10 (5%) within gene exons, 134 (66%) within gene introns and 1 (<1%) within 1 kb upstream of annotated genes.

A total of 204 integration sites were identified in the C+V group and 179 in the C then V group, distributed across all chromosomes (Figure 7). Five shared sequence variants were identified in both samples, suggesting these originated from the viral construct itself rather than de novo integrations. Integration site annotation revealed that in the C+V sample, 10 (5%) insertions occurred in exons, 134 (66%) in introns, and 1 (<1%) within 1 kb upstream of annotated genes (Figure 7D). Similarly, in the C then V sample, 9 (5%) were found in exons, 124 (69%) in introns, and 2 (1%) within 1 kb upstream of genes (Figure 7D). Thus, the overall integration landscape remained comparable between groups, demonstrating that CRISPR-induced DSBs at *TRAC* and *TRBC* do not elevate transgene TCR integration at these sites. The majority of integration sites occurred within introns, likely without direct effects on gene expression. However, as approximately 5% of insertions were in exon regions, there remains a small theoretical risk integration could disrupt oncogenes or tumor suppressor genes. These findings underscore the importance of comprehensive genomic safety screening for lentivirally-engineered T cells intended for clinical application.

### CRISPR-mediated removal of endogenous TCRs enhances expression and function of autoreactive TCRs

To determine whether the benefits of endogenous TCR elimination extend to autoreactive TCRs, we evaluated four insulin-reactive TCRs: 1C8 (HLA-A2/PPI:2–12)^44^, 1E6 (HLA-A2/PPI:15–24)^45, 46^, GSE.20D11 (HLA-DQ8/InsB:9–23)^47, 48^, and Clone 5 (HLA-DQ8/InsB:9–23)^49-52^. HIS mice were generated by reconstituting thymectomized NSG hosts with HLA-A2⁺ CD34⁺ human HSCs and autologous human fetal thymus. At 16 weeks post-transplantation, a reconstituted mouse in the cohort was euthanized and its splenic T cells were transduced with these four autoreactive TCRs with and without CRISPR-mediated removal of endogenous TCRs. In all cases, transduction efficiency was higher in T cells receiving combined CRISPR plus lentiviral transduction (C+V) compared to lentiviral transduction alone (V) (Figure 8A). Similar to our MART-1 studies, removal of endogenous TCRs also reduced the generation of mispaired/unwanted TCRs, as reflected by a decrease in dextramer⁻ (or Vβ⁻) populations within the reporter⁺ cells. We next focused on the HLA-A2–restricted class I TCRs 1C8 and 1E6. T cells transduced with these receptors (V vs. C+V) were co-cultured with SC-islets that lacked all HLA except HLA-A2^53^ (referred to henceforth as “HLA-A2–only”), which removed the possibility of alloresponses to non-A2 HLAs. As T cells in these HIS mice were generated in an HLA-A2⁺ fetal thymus and from HLA-A2⁺ HSCs, they are expected to be tolerant to HLA-A2 and thus should not mount an alloresponse to HLA-A2–only SC-islets. After 48 hours of co-culture, 1E6-transduced T cells in the C+V group showed a significantly higher activation index (%CD69⁺ in reporter⁺ vs reporter⁻ populations) than their V group counterparts, whereas 1C8 TCR showed no significant difference (Figure 8B).

**Figure 8.**
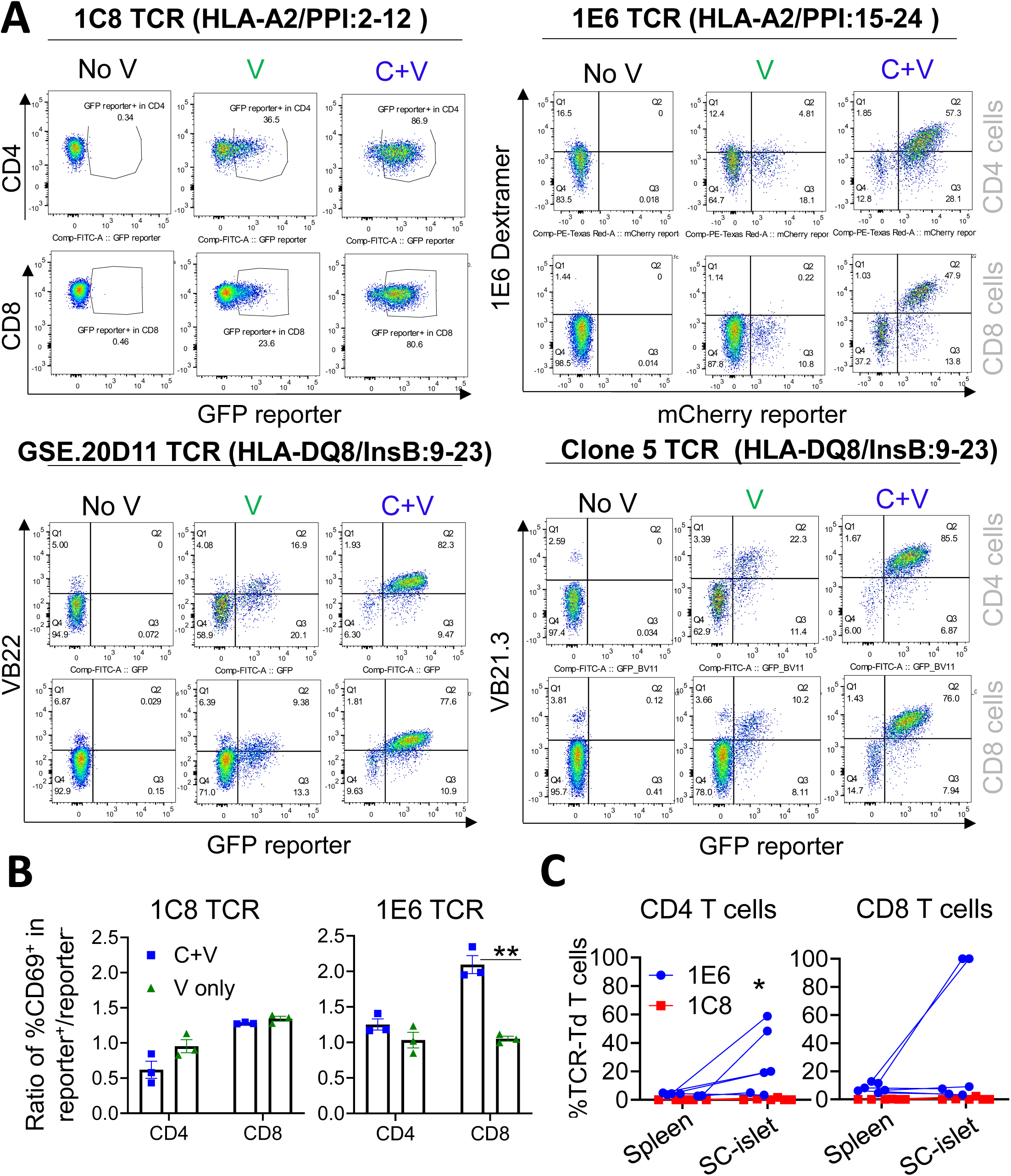
CRISPR-mediated removal of endogenous TCRs enhances expression and function of autoreactive TCRs. (A) Transduction efficiency of four insulin-reactive TCRs—1C8 (HLA-A2/PPI:2–12), 1E6 (HLA-A2/PPI:15–24), GSE.20D11 (HLA-DQ8/InsB:9–23), and Clone 5 (HLA-DQ8/InsB:9–23)—was compared between lentivirus-only (V) and CRISPR plus lentivirus (C+V) conditions. Transduction efficiency and mispairing frequency (reporter⁺ dextramer⁻ or Vβ⁻) were quantified for each TCR. (B) Functional activity of class I TCRs 1C8 and 1E6 was assessed by co-culture of TCR-transduced T cells (V vs C+V) with HLA-A2–only SC-islet cells. The activation index was calculated as the ratio of %CD69⁺ cells in reporter^+^ versus reporter⁻ populations. (C) *In vivo* function was evaluated in HIS mice bearing thigh muscle grafts of HLA-A2–only SC-islets. A mixed population of TCR-transduced T cells (C+V condition) was adoptively transferred. At endpoint, 1E6 TCR-transduced T cells were significantly enriched within SC-islet grafts compared with spleens. Data in panels (B) and (C) were analyzed by unpaired and paired t tests, respectively (* 0.01<p-value<0.05, **0.001<p-value<0.01).

To further assess *in vivo* functionality, SC-islets were engrafted into the thigh muscle of HIS mice at week 10 post-humanization. At week 18, mice received intravenous injections of C+V-edited T cells transduced with either 1E6 and 1C8 (n=3) or all four TCRs (1E6, 1C8, GSE.20D11, and Clone 5) (n=3). Four weeks post-transfer, 1E6 TCR-transduced T cells were significantly enriched within SC-islet grafts compared with spleens (Figure 8C). In contrast, GSE.20D11 and Clone 5 T cells were undetectable in either grafts or spleens, consistent with the absence of HLA-DQ8 in both SC-islets and HSCs.

Histological analysis of SC-islet grafts from mice receiving TCR-transduced T cells (n = 6) revealed islet destruction characterised by dense immune infiltrates, collagen deposition, and disrupted architecture. As a positive control for alloimmunity, HLA-A2–only SC-islet grafts transplanted into HIS mice reconstituted with HLA-A2⁻ thymus and HSCs also showed marked inflammatory destruction (n = 3). In contrast, grafts from a non-humanized control and a HIS mouse reconstituted with HLA-A2⁺ thymus/HSCs but lacking TCR-transduced T cells displayed intact islet structure with no evidence of infiltration (Figure S6A). Imaging mass cytometry confirmed infiltration of CD4⁺ and CD8⁺ T cells in the SC-islet grafts (Figure S6B). Together, these findings demonstrate that CRISPR-mediated removal of endogenous TCRs enhances both the expression and functional activity of autoreactive TCRs. This highlights the broader applicability of this strategy for dissecting autoimmune mechanisms and for engineering precision TCR-based therapies targeting self-antigens.

## Discussion

Adoptive transfer of genetically engineered T cells has transformed cancer immunotherapy, providing the means to redirect immune specificity to target and destroy tumor cells ^2, 54^. Although CAR-T cells have demonstrated remarkable success in B-cell malignancies^55^, their restriction to surface antigens limits their reach. In contrast, TCR-based approaches can target of intracellular and neoantigens enabling treatment of solid tumors^56^. However, therapeutic translation of TCR engineering has been hindered by competition with endogenous receptors and the risk of mispaired, potentially autoreactive heterodimers of transduced and endogenous TCR chains^18, 19^. Eliminating endogenous TCRs is therefore central to maximizing both safety and efficacy.

Here we describe a universal CRISPR/Cas9 strategy that achieves complete and selective knockout of endogenous TCR-α and -β chains without affecting transduced TCRs irrespective of codon optimization. Dual chain targeting provides a general solution to TCR mispairing and establishes a single-step manufacturing platform that is faster and more reproducible than current multistage protocols. The approach not only simplifies production but also enhances transgene expression and antigen sensitivity, yielding T cells with superior potency *in vitro* and *in vivo*.

By simultaneously delivering Cas9-gRNA ribonucleoproteins and lentiviral vectors (C+V), we achieved efficient deletion and transduction in the same electroporation step, removing the need for retronectin and extended centrifugation. Functionally, endogenous TCR deletion increased DMF5 TCR surface expression, boosted cytokine release, and augmented tumor-specific cytotoxicity. In HIS mice, a model that most closely recapitulates human T-cell development and immune interactions *in vivo*, these edited cells mediated robust tumor regression with durable persistence and no evidence of graft-versus-host disease (GvHD), representing, to our knowledge, the first demonstration of CRISPR-edited, mispair-free human TCR function in a humanised in vivo system.

Our integration site analysis provides further reassurance regarding safety. Despite CRISPR-induced double-strand breaks at *TRAC* and *TRBC* loci, lentiviral insertions remained evenly distributed across the genome with no enrichment at cut sites. This confirms that genome editing did not bias viral integration and supports the genomic stability of the C+V approach. Although a small proportion of integrations occurred in exonic regions, this frequency is consistent with prior reports for lentiviral vectors used in approved CAR T cell products^57, 58^. The approach could be refined further by combining CRISPR editing with integrase-deficient lentiviruses to insert transgenes into specific sites through HDR^59^ or non-viral approaches relying on HDR CRISPR for site-specific transgene insertion^28, 30, 60^.

Importantly, the same CRISPR workflow that improves tumor-specific TCR function also enables clean, mispair-free expression of autoreactive TCRs. By applying the method to four insulin-reactive TCRs, we demonstrated that deletion of endogenous TCRs markedly enhanced transduction efficiency and restored functional activity even for low-affinity autoimmune receptors. The ability to express and test human autoreactive TCRs in a defined, non-mispairing background provides a unique experimental system for dissecting autoimmune pathogenesis and tolerance mechanisms. The 1E6 TCR^45, 46^ in particular showed strong activation against stem cell-derived HLA-A2–only islets *in vitro* and infiltration of grafts *in vivo*. These findings validate the platform as a model of human autoimmune disease and point toward a next generation of humanized mouse models in which human insulin and HLA transgenes replace their murine counterparts, creating an almost fully human immune– target interface for studying autoimmune mechanisms and testing therapeutic interventions.

In summary, simultaneous CRISPR deletion of endogenous TCR-α and -β chains provides a universal, streamlined, and clinically adaptable platform for TCR engineering. By preventing mispairing, enhancing potency, and enabling faithful *in vivo* modelling of both tumour immunity and autoimmunity in human immune system mice, this approach bridges fundamental immunology and translational therapy within a single experimental framework. It establishes a foundation for developing safer, more precise TCR-based immunotherapies and next-generation humanized models of immune pathology.

## Materials and Methods

### sex as a biological variable

Our study examined male and female animals, and similar findings are reported for both sexes.

### Lentiviral vector production

A second-generation lentiviral system (pHR-EF1α_IRES_GFP_SIN backbone) was used to produce viral supernatants carrying MART-1 TCR (clone DMF5)^61^, Clone 5 TCR^49-52^, 1E6 TCR^44-46^, 1C8 TCR^44^ and GSE.20D11 TCR^47, 48^ or the MART-1 peptide (H3 construct on pLVX backbone)^61^. HEK293T were transfected with Lipofectamine 2000 (Thermo Fisher). Viral supernatants were concentrated by ultracentrifugation and stored at -80°C until use.

### Cell lines and culture

K562-HLA-A2 cells were kindly provided by Dr. James Riley (University of Pennsylvania), and T2 cells were obtained from ATCC. PBMCs from eight anonymous healthy donors were sourced from the New York Blood Center and isolated by Histopaque (Sigma Aldrich) density centrifugation. B lymphoblastoid cell lines (LCLs) were generated by EBV infection of magnetic-activated cell sorting (MACS)-sorted B cells from HLA-A2^+^ PBMCs and cultured for 3 months ^62^. Rremaining PBMCs were cryopreserved for autologous T-cell transduction. On day 0, thawed PBMCs were stimulated with ImmunoCult™ Human CD3/CD28/CD2 Activator (25 µL/mL, StemCell Technologies) in X Vivo 15 media (Lonza) supplemented with 10% human serum (Gemini Bio-Products), 100 IU/mL IL-2, 10 ng/mL IL-7, and 10 ng/mL IL-15.

### T-cell transduction

Retronectin-coated 96-well plates (25 μg/mL, Clontech) were loaded with lentivirus (MOI of 10-20) and centrifuged at 1500 x g for 90 min at 32°C. Activated T cells were then added and centrifuged at 500 x g for 10 min at 32°C before overnight incubation. The following day, medium was replaced with cytokine-supplemented media (IL-2, IL-7, IL-15) and cells were expanded with irradiated allogeneic PBMCs and LCLs plus PHA.

### CRISPR-mediated TCR removal

TCR removal was achieved by electroporating Cas9 RNP complexes on day 2 post-activation, either simultaneously with TCR lentivirus (in the C+V group) or 1 h prior (C then V group). crRNAs (Table S1, IDT) targeting *TRAC* and *TRBC* were annealed with tracrRNA and complexed with recombinant Cas9 (IDT). RNPs were electroporated into 2-3x10^5^ T cells using the Neon Transfection System (Thermo Fisher) at 1600 V voltage, 10 ms width, and 3 pulses.

### Human pluripotent stem cell culture and SC-islet differentiation

NIH-approved INSGFP/W hESCs (MEL-1, NIH #0139) were cultured on irradiated mouse embryonic fibroblasts in standard maintenance media and differentiated into stem cell–derived islets (SC-islets) following established protocols ^63, 64^. Full differentiation media compositions are provided in Supplementary Methods^65^. G-banded karyotyping confirmed genomic stability before use.

### Human fetal tissues

All human fetal liver and thymus (17–21 weeks gestation) were obtained from Advanced Bioscience Resources (Alameda, CA) under approved protocols. Fetal liver CD34^+^ hematopoietic stem cells were isolated using MACS positive selection (Miltenyi Biotec) as described^66^.

### Mouse studies

NSG (NOD.Cg-*Prkdc^scid^ Il2rg^tm1Wjl^*/SzJ) and NSG-MHC I/II DKO (NOD.Cg-*Prkdc^scid^ H2-K1^tm1Bpe^ H2-Ab1^em1Mvw^ H2-D1^tm1Bpe^ Il2rg^tm1Wjl^*/SzJ) mice (Jackson Laboratory) were bred or maintained under pathogen-free conditions. All procedures were approved by the Columbia University Institutional Animal Care and Use Committee. Mice were thymectomized^67^, sublethally irradiated and reconstituted with 1.5-2×10^5^ human CD34^+^ fetal liver cells to generate human immune system (HIS) mice. This approach provided a uniquely humanized in vivo platform to assess TCR-engineered T cells. For tumor studies, 7-10-week-old NSG-MHC I/II DKO HIS mice received intradermal injections of 1×10^6^ K562 HLA-A2^+^ cells expressing or lacking the MART-1 peptide in Matrigel (Corning) on the right and left flanks, respectively, followed by intravenous injection of 2 × 10^6^ T cells on day 3.

For autoimmunity studies, HIS mice were reconstituted with HLA-A2^+^ or HLA-A2^-^ HSCs and autologous (to HSCs) fetal thymus fragments. At week 10 post ∼3000 HLA-A2–only SC-islet clusters were engrafted into thigh muscle. At week 18 post-reconstitution, one mouse in the cohort with HLA-A2^+^ HSCs was euthanized and splenic T cells were activated (for 48 hours) and transduced with the 4 insulin-specific TCRs (1E6, 1C8, Clone 5, and GSE.20D11). After two weeks of *ex vivo* expansion of TCR-transduced T cells (around week 18 post humanization), other mice in the cohort were adoptively transferred (4×10^6^ T cells per TCR per mouse) with a mixture of T-cells transduced with 1E6, 1C8, Clone 5 and GSE.20D11 TCR (C+V condition) (3 mice with all 4 TCRs and 3 mice with 1E6+1C8 only). Mice were euthanized four weeks later (about week 22 post-reconstitution) and analyzed by flow cytometry and imaging mass cytometry.

### Histological Imaging

Tissues were fixed in 4% formaldehyde for 24-48 hours, paraffin-embedded, sectioned (5 µm) sections and stained with H&E^68^. Slides were scanned (Leica SCN 400) analyzed using HALO (Indica Labs) ^69, 70^. For imaging mass cytometry, formalin-fixed SC-islet grafts were processed and stained with a 37-antibody panel as described^71^. Acquisition was performed using the Hyperion Imaging System (Standard BioTools) at 200 Hz, and images were analyzed in HALO v4.1. Further details on imaging mass cytometry are provided in the Supplementary Methods.

### Flow cytometry

Cells were stained with fluorochrome-conjugated antibodies listed in Table S3. For intracellular staining, the Foxp3/Transcription Factor Buffer Set (Thermo Fisher) was used. Blood samples (50 µL) were processed with ACK lysis buffer, and counting beads were added for quantification. Samples were acquired on Fortessa (BD Biosciences) or Aurora (Cytek) instruments. Data analyses were performed using FlowJo software (Tree Star).

### Targeted Locus Amplification

TLA sequencing was performed by Cergentis (Utrecht, Netherlands) as described^43, 72^, using primers (Table S2) targeting the transgenic RRE or GFP regions (Table S2). Integration sites were mapped to the human genome (GRCh38) to assess transgene distribution and proximity to *TRAC*/*TRBC* loci. Further details on TLA sequencing are provided in the Supplementary Methods.

### Statistical Analysis

Data were analyzed using Graphpad Prism v8.0 or v9.0. Statistical significance was determined by Student’s T tests or ANOVA with p < 0.05 were considered significant. Data are presented as mean ± SEM.

## Data Availability Statement

All data supporting the findings are included within the article and its supplementary information files. The raw data that support the findings of this study are available from the corresponding author upon reasonable request.

## Acknowledgments

This work was supported by R21AI175813 (Khosravi-Maharlooei), R01AI177872, P01AI045897 (both Sykes). Translational Therapeutics (TRx) and Biomedical Engineering Technology Accelerator (BiomedX) Pilot Award (Khosravi-Maharlooei), Nelson Foundation Faculty Development Award (Khosravi-Maharlooei), R01AI177872 (Sykes), P01AI045897 (Sykes), Juvenile Diabetes Research Foundation 2-SRA-2022-1220-S-B (Sykes and Creusot), 3-PDF-2020-944-A-N (Teteloshvili), R01AI142428 (Creusot) and the NIDDK-supported Human Islet Research Network (HIRN) (U01DK123559) (Sykes). MKM was previously supported by an American Diabetes Association (ADA) Postdoctoral Fellowship and a Columbia University Naomi Berrie Diabetes Center Russell Berrie Foundation Fellowship. Some of the studies were conducted at the Columbia Center for Translational Immunology (CCTI) Flow Cytometry Core facility, which is supported in part by the grants S10OD020056 (Influx sorter), S10RR027050 (LSR-II), P30CA013696 (Herbert Irving Cancer Center) and 5P30DK063608 (Diabetes Research Center). AKS is a Wellcome Investigator (220295/Z/20/Z). The content is solely the responsibility of the authors and does not necessarily represent the official views of the NIH. We thank Ms. Julissa Cabrera (Columbia University) for assistance with the submission.

## Author contributions

Conceptualization: MKM, ML, RJC, GZ, and MS; methodology: MKM, ML, RJC, DT, REH, AVP, GZ, and MS; software and formal analysis: GZ, AC, NS and MKM; validation: MKM and MS; investigation: MKM, GZ, HL, AC, FF, NT and XD; resources: RJC and AKS; writing original draft: GZ, NS and MKM; writing review and editing: MKM, RJC, AVP, KHK, EB, AKS, and MS; funding acquisition: MS, and MKM; supervision: MKM.

