## Supplement for "A universal platform for simultaneous TCRα/β removal enables safer and more potent TCR therapies and autoimmune modeling"

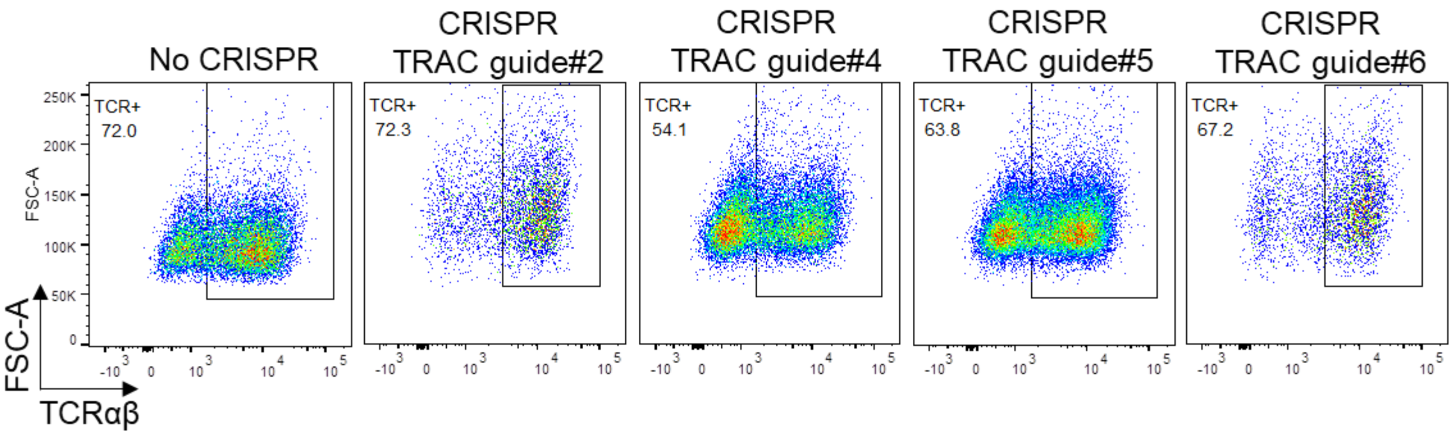


**Figure S1. Efficiency of TCRαβ knockout with different sub-optimal guide RNAs targeting TCRα chain.** Jurkat cells were electroporated with ribonucleoprotein complexes containing Cas9 protein and different guide RNAs targeting TCRα chain. After 72h, the knockout efficiency was measured through flow cytometry staining of TCRαβ. Results for effective guides (TRAC1 and TRAC3) are shown in Figure 1B. Sequences of all the guides targeting TCR α and β chains are shown in Table S1.

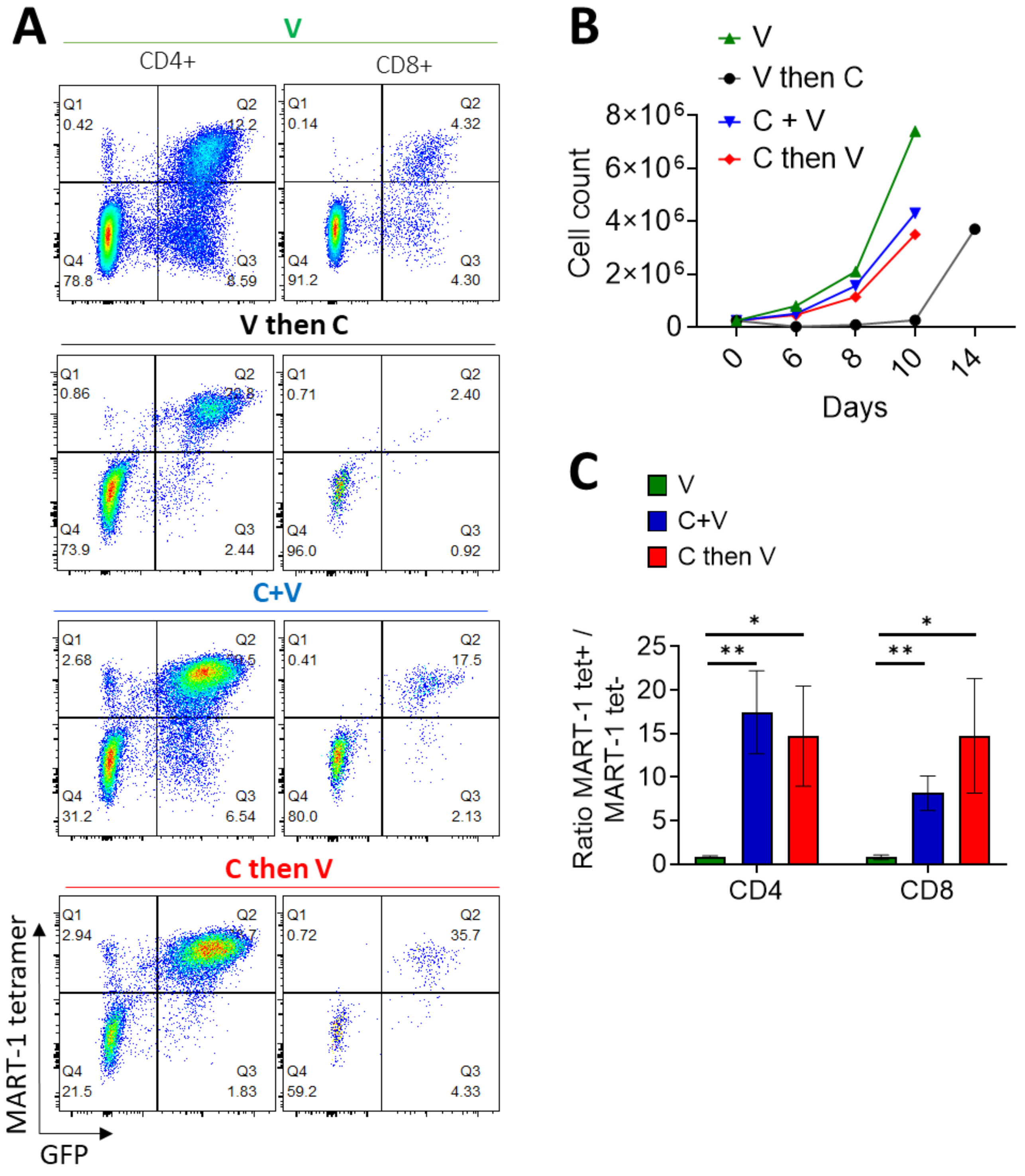


**Figure S2. Comparing different strategies for combined removal of endogenous TCRs and transduction of transgenic TCRs.** (A) Expression of DMF5 TCR (tetramer staining) and reporter gene (GFP) was measured in different groups. (B) Expansion of engineered T cells in different groups was compared up to 14 days after initial activation. (C) Ratio of MART-1 tetramer-positive to tetramer-negative cells within the transduced T cells (GFP$^{+}$).

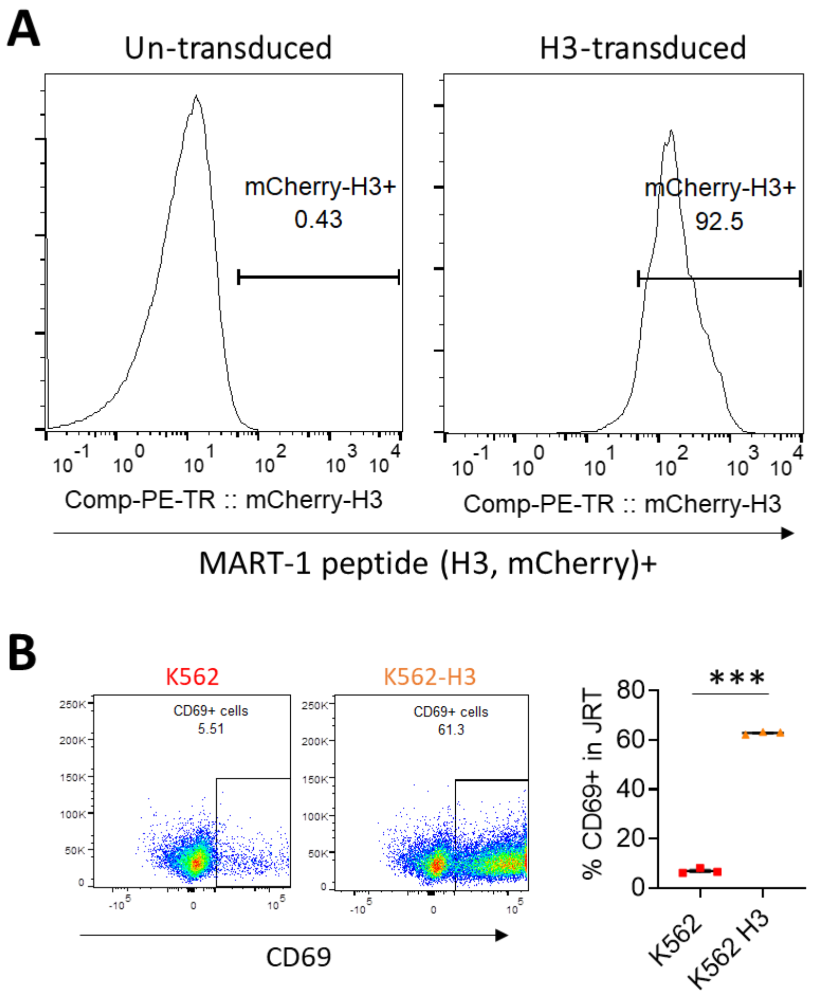


**Figure S3. Transduction of HLA-A2⁺ K562 cells with MART-1 peptide (H3) and antigen-specific activation of DMF5 TCR.** (A) Transduction efficiency in HLA-A2⁺ K562 cells was quantified by flow cytometry using mCherry reporter positivity. (B) To assess antigen-specific activation, MART-1–peptide–expressing K562 cells (K562-H3) were co-cultured for 24 h with TCR-deficient Jurkat (JRT3.1) cells engineered to express the DMF5 TCR. DMF5 TCR⁺ JRT3.1 cells upregulated CD69 following exposure to K562-H3, confirming DMF5 TCR activation.

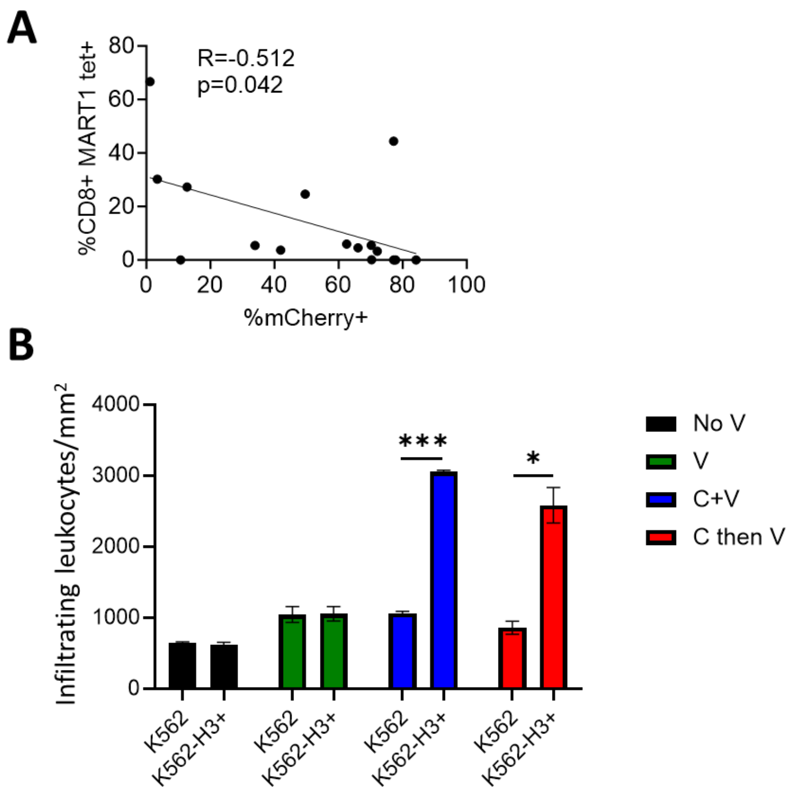


**Figure S4. Reverse correlation between the number of K562 HLA-A2⁺ MART-1 peptide⁺ mCherry⁺ tumor cells and the number of MART1 tetramer⁺ CD8⁺ infiltrating T cells present in tumors explanted from recipient HIS mice (see Figure 4E).** (A) As the frequency of CD8⁺ MART-1⁺ T cells increases, the number of remaining K562 HLA-A2⁺ MART-1 peptide⁺ mCherry⁺ cells decrease. (B) Quantification of tumor infiltrating leukocytes in histologic images (H&E staining) of K562 HLA-A2⁺ and K562 HLA-A2⁺ MART-1 peptide⁺ tumors treated under various conditions (see Figure 5) (n=3 mice/group).

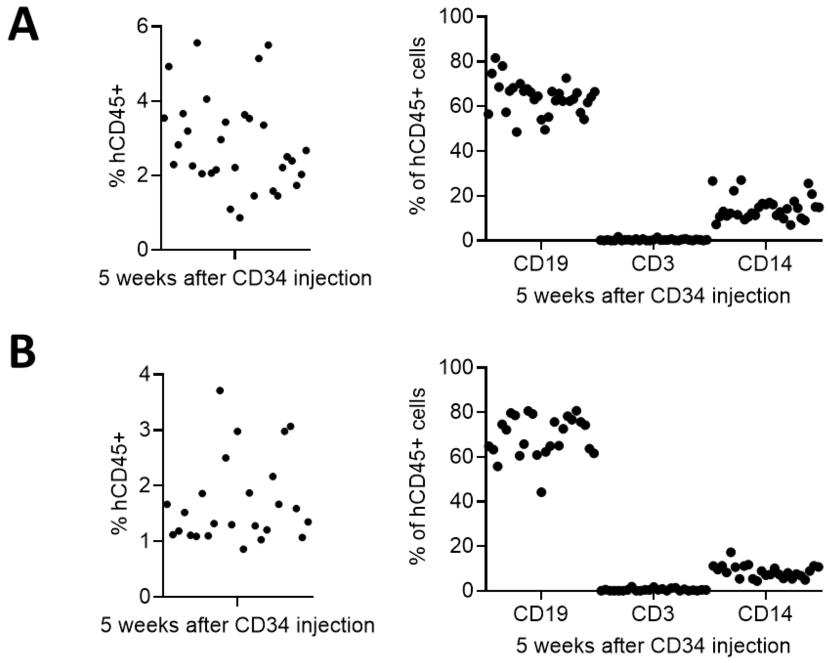


**Figure S5. Human cell reconstitution of mice that have been thymectomized and injected with fetal liver CD34$^+$ cells.** (A) Levels of human hematopoietic cells (hCD45) including human B cells (CD19), T cells (hCD3), and monocytes (hCD14) in the blood of HIS mice used for the tumor study (see Figure 4C). (B) Levels of human hematopoietic cells (hCD45) including human B cells (CD19), T cells (hCD3), and monocytes (hCD14) in the blood of the HIS mice used for the GvHD study (see Figure 6A).

**A**

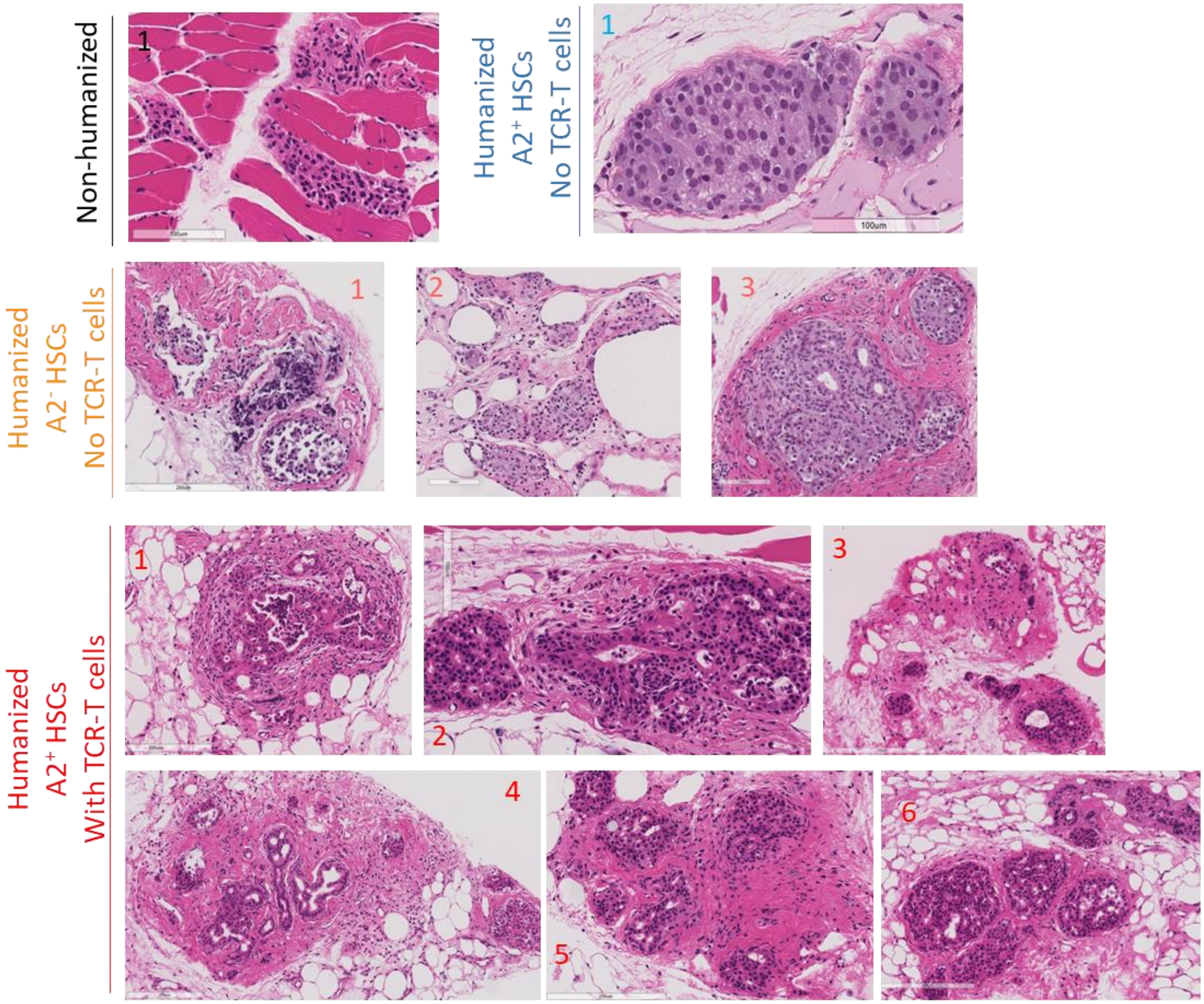

Non-humanized
1
Humanized
A2+ HSCs
No TCR-T cells
1
100um
Humanized
A2- HSCs
No TCR-T cells
1
2
3
Humanized
A2+ HSCs
With TCR-T cells
1
2
3
4
5
6


**B**

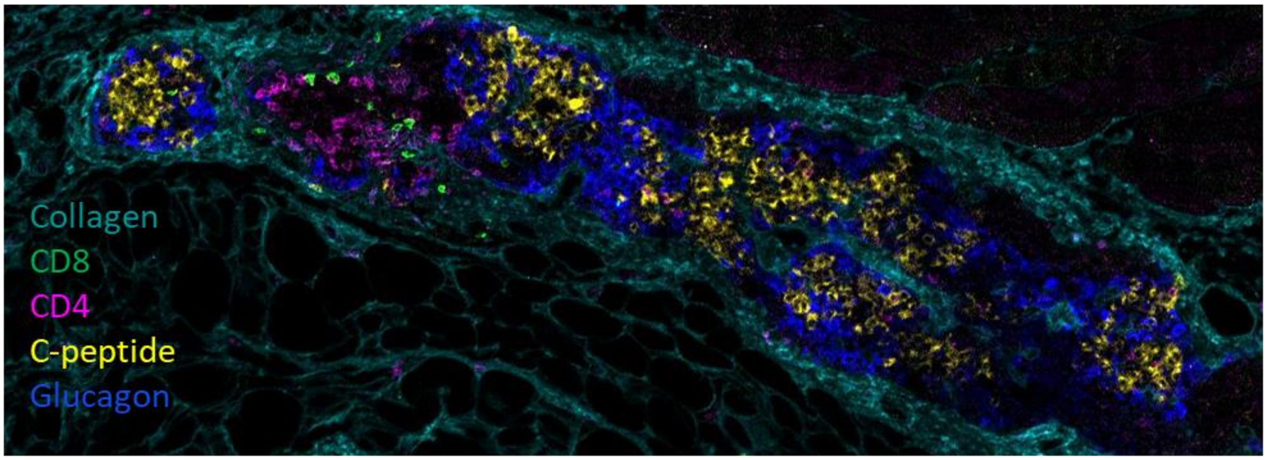

Collagen
CD8
CD4
C-peptide
Glucagon

**Figure S6. Histopathology and immune profiling of SC-islet grafts in HIS mice.** (A) H&E staining of SC-islet grafts from mice receiving TCR-transduced T cells (n=6) shows graft destruction characterized by dense immune infiltrates, collagen deposition, and disrupted islet architecture. As an alloimmunity control, HLA-A2–only SC-islet grafts transplanted into HIS mice generated with HLA-A2$^{-}$ HSCs and thymus (n=3) also exhibit destructive inflammatory infiltrates. In contrast, grafts in a non-humanized mouse and in a control HIS mouse reconstituted with HLA-A2$^{+}$ thymus/HSCs but lacking TCR-transduced T cells show no histopathologic evidence of injury or infiltration. (B) Imaging mass cytometry shows robust infiltration of CD3$^{+}$, CD4$^{+}$, and CD8$^{+}$ T cells in SC-islet graft of a mouse that received TCR-transduced T cells, with immune cells enriched around C-peptide$^{+}$ and glucagon$^{+}$ islet regions.

**Table S1:** Sequence of guide RNAs targeting TRAC and TRBC intron/exon boundaries as indicated in Figure 1A.

| TCR gene | Guide | Sequence |
|---|---|---|
| TRBC1+TRBC2 | 1 | CCTGGGGTAGAGCAGGTGAG |
| TRBC1+TRBC2 | 2 | CCACTCACCTGCTCTACCCC |
| TRAC | 1 | CAGGGTTCTGGATATCTGT |
| TRAC | 2 | CTGCCCTTACCTGGGCT |
| TRAC | 3 | CATCACAGGAACTTTCTAAA |
| TRAC | 4 | AGCTTTGAAACAGGTAAGAC |
| TRAC | 5 | TTCGTATCTGTAAAACCAAG |
| TRAC | 6 | TCAAGGCCCCTCACCTCAGC |

**Table S2:** Sets of primers used to perform TLA.

| Primer | Sequence |
|---|---|
| RRE Rv | TTCACTTCTCCAATTGTCCC |
| RRE Fw | TGGCTGTGGAAAGATACCTA |
| GFP Rv | GTCATCCTCGAAGAAGATGG |
| GFP Fw | TGTACATCATGACCGACAAG |

**Table S3:** List of flow cytometry antibodies used in the study.

| Marker | Fluorophore | Clone and Vendor |
|---|---|---|
| CD8 | APC-Cy7 | SK1, BD Pharmingen |
| CD4 | AF700 | OKT4, Tonbo |
| CD3 | BV650 | UCHT1, BD Pharmingen |
| CD45 | AF660 | HI30, BioLegend |
| CD69 | BV650 | FN50, BioLegend |
| CD25 | BV711 | M-A251, BioLegend |
| CD19 | BV510 | HIB19, BioLegend |
| IFN-ɣ | APC | 4S.B3, BioLegend |
| TCRαβ | PE | IP26, BioLegend |
| Ki-67 | APC | Ki-67, BioLegend |
| HLA-DR | BV605 | G46-6, BD Pharmingen |
| HLA-A*02:01 Mart-1 tetramer (ELAGIGILTV) | PE | SFCI21Thy2D3, MBL |

Supplementary Methods.

*Human pluripotent stem cell culture and SC-islet differentiation*

Pluripotent cells were maintained on irradiated mouse embryonic fibroblasts (Thermo Fisher) in hESC maintenance media composed of Dulbecco's Modified Eagle Medium (DMEM)/F12, 20% (vol/vol) Knockout serum replacement (Thermo Fisher Scientific), nonessential amino acids (Thermo Fisher Scientific), GlutaMAX (Thermo Fisher Scientific), and beta-mercaptoethanol (Millipore). The maintenance media was supplemented with 10 ng/mL recombinant human FGF-2 (R&D Systems). Confluent hESCs were dissociated into single-cell suspension with TrypLE Select (Gibco) and passaged every 3–4 days. G-banded karyotyping performed by Cell Line Genetics confirmed normal karyotype of INSGFP/W hESCs.

To initiate differentiation, confluent cultures were dissociated into single-cell suspensions using TrypLE Select, counted, and seeded in six-well suspension plates at a density of 5.5 x $10^6$ cells per 5.5 mL of hESC maintenance media supplemented with 10 ng/mL activin A (R&D Systems) and 10 ng/mL heregulin B (PeproTech). The plates were incubated at 37°C with 5% $CO_2$ on an orbital shaker set at 100 rpm to induce 3D sphere formation. After 24 hours, the spheres were collected and allowed to settle by gravity, washed once with RPMI medium (Gibco), and resuspended in day 1 differentiation media. The resuspended spheres were distributed into fresh six-well suspension plates for a final volume of 5.5 mL of day 1 media per well. Until day 3, spheres were fed daily by removing 5 mL of the media and replenishing with 5.5 mL. From days 4 to 20, 5 mL media was removed daily and 5 mL of fresh media was added. Media compositions for differentiation are as follows: day 1, RPMI (Gibco) containing 0.2% fetal bovine serum (FBS, Corning), 1:5000 insulin transferrin selenium G (ITS-G, Gibco), 100 ng/mL activin A (R&D Systems), and 50 ng/mL WNT3a (R&D Systems); day 2, RPMI containing 0.2% FBS, 1:2000 ITS, and 100 ng/mL activin A; day 3, RPMI containing 0.2% FBS, 1:1000 ITS, 2.5 μM TGFbi IV (Calbiochem), and 25 ng/mL keratinocyte growth factor (KGF; R&D Systems); day 4–5, RPMI containing 0.2% FBS, 1:1000 ITS, and 25 ng/mL KGF; day 6–7, DMEM (Gibco) with 25 mM glucose containing 1:100 B27 (Gibco) and 3 nM TTNPB (Sigma); day 8, DMEM with 25 mM glucose containing 1:100 B27, 3 nM TTNPB, and 50 ng/mL epidermal growth factor (EGF; R&D Systems); day 9–11, DMEM with 25 mM glucose containing 1:100 B27, 50 ng/mL EGF, and 50 ng/mL KGF; day 12-19, DMEM with 25 mM glucose containing 1:100 B27, 1:100 Gluta-MAX (Gibco), 1:100 NEAA (Gibco), 10 μM ALKi II (Axxora), 500 nM LDN-193189 (Stemgent), 1 μM Xxi (Millipore), 1 μM T3 (Sigma-Aldrich), 0.5 mM vitamin C, 1 mM N-acetylcysteine (Sigma-Aldrich), 10 μM zinc sulfate (Sigma-

Aldrich), and 10 µg/mL heparin sulfate; day 20+, Connaught Medical Research Laboratories (CMRL) medium containing 10% FBS, 1:100 Glutamax (Gibco), 1:100 NEAA (Gibco), 10 µm ALKi II (Axxora), 0.5 mM vitamin C, 1 µM T3 (Sigma-Aldrich), 1 mM N-acetyl Cysteine (Sigma-Aldrich), 10 µM zinc sulfate (Sigma-Aldrich), and 10 µg/mL of heparin sulfate. The cells were also supplemented with 1 mM aphidicolin (Cayman Chemical) starting on day 13.

*Imaging mass cytometry analysis*

At the time of sacrifice, ES-derived HLA-A2–only SC-islet grafts were fixed in formalin 10% and then transferred to ethanol for histological and IMC analysis. IMC was performed on sections of formalin-fixed, paraffin-embedded samples. Briefly, tissue sections were cut at 6 µm and mounted on slides, which were then deparaffinized in 2 x 20 min washes of xylene and then gradually rehydrated by sequential washes from 100% through 70% ethanol. Slides were then washed briefly in PBS and transferred to Tris/EDTA (10 mM Tris, 1 mM EDTA, pH 9.2) buffer for antigen retrieval in a decloaking chamber (Biocare Medical) at 95°C for 30 minutes before being cooled at RT for 1 hour. Slides were then blocked in 3% BSA in PBS for 1 hour at RT and stained with ~100 µL/slide of the antibody cocktail overnight at 4°C in a humidified chamber. The next day, slides were stained for DNA by incubating for 30 minutes at RT with 1.25 µM Cell-ID Intercalator-Ir (Standard Biotools). Antibody and DNA-staining cocktails were diluted in 0.5% BSA in PBS. Batches of slides were then washed one time in an excess (~200 mL) of PBS and then twice in ultrapure 18.2 MΩ water, before being air dried and acquired on the IMC (Standard BioTools, Hyperion Imaging System) according to Standard Biotools's standard operating procedures, with a 200 Hz laser frequency and a 1 µm step increment. The final panel consisted of 37 antibodies. Images were analyzed using Halo software (Indica Labs, version 4.1).

*Targeted Locus Amplification*

TLA sequencing was performed by Cergentis (Utrecht, Netherlands). Briefly, DNA was crosslinked, fragmented, re-ligated, and de-crosslinked. This product served as the TLA template, which was subsequently fragmented, circularized, and amplified with inverse primers complementary to a short locus-specific sequence. Once the complete locus was amplified, ~2 kb segments were sheared. Libraries were prepared for sequencing on an Illumina platform. Two sets of primers (Table S2) targeting the transgenic RRE or GFP sequences regions were used in individual TLA amplifications. Integration sites were mapped to the human genome (GRCh38) to assess transgene distribution and proximity to *TRAC*/*TRBC* loci.
